# Paracrine granules are cytoplasmic RNP granules distinct from stress granules that assemble in response to viral infection

**DOI:** 10.1101/2021.08.06.455464

**Authors:** Valentina Iadevaia, James M. Burke, Lucy Eke, Carla Moller-Levet, Roy Parker, Nicolas Locker

## Abstract

To rapidly respond and adapt to stresses, such as viral infections, cells have evolved several mechanisms, which include the activation of stress response pathways and the innate immune response. These stress responses result in the rapid inhibition of translation and condensation of stalled mRNAs, together with RNA-binding proteins and signalling components, into cytoplasmic biocondensates called stress granules. Increasing evidence suggests that stress granules contribute to antiviral defense and thus viruses need to evade these response pathways to propagate. In addition, the stress granule pathway is proposed to be dynamic and adaptable to specific stresses. We previously showed that Feline Calicivirus (FCV) impairs SGs assembly by cleaving the scaffolding protein G3BP1. We also observed that uninfected bystander cells assembled G3BP1-granules, suggesting a paracrine response trigged by the infection. We now present evidence that virus-free supernatant generated from infected cells can induce the formation of paracrine granules. They are different from canonical stress granules and exhibit specific kinetics of assembly-disassembly, protein and RNA composition and are linked to antiviral activity. We propose that this paracrine induction reflects a novel cellular defence mechanism to limit viral propagation and promote stress responses in bystander cells.

**Summary statement:** We describe a novel type of paracrine induced RNA granules associated with viruses, highlighting how different stresses results in heterogeneous stress granule-like condensates with specific cellular functions.

## Introduction

Controlling the localisation and function of macromolecules is central to cell biology and can be achieved by surrounding them with lipid membranes in organelles such as the nucleus, lysosomes or mitochondria. Membraneless compartments, known as biocondensates or membraneless organelles, are increasingly recognised as an alternative way to organise cellular components. They are maintained through a combination of protein-protein, protein-RNA and RNA-RNA interactions and their dynamic formation generates high local concentrations of RNA and protein (Tauber et al., 2020). Because of this remarkable molecular plasticity, biocondensates provide an ideal platform for the regulation of cellular fundamental processes, such as mRNA metabolism or intracellular signalling, and utilised by cells to rapidly adjust and rewire regulatory networks in response to various physiological and pathological triggers (Yoo et al., 2019).

Stress granules (SGs) are among the most characterised cytoplasmic biocondensates (Corbet and Parker, 2019; Hofmann et al., 2020). They are important for the organisation of cellular content, capturing mRNAs and proteins during stresses including oxidative stress, heat shock, viral infection, proteasomal inhibition, ER stress, UV irradiation, among others (Hofmann et al., 2020). The general inhibition of protein synthesis following stress, usually triggered by phosphorylation of the eukaryotic translation initiation factor 2α (eIF2α), results in the dissociation of mRNAs from polysomes and their accumulation in RNP complexes (Hofmann et al., 2020). This increased concentration of cytoplasmic RNPs and their binding by aggregation prone RNA-binding proteins (RBPs), such as Ras-GTPase activating SH3 domain binding protein 1 (G3BP1) and T cell internal antigen-1 (TIA-1), results the recruitment of multiple proteins characterised by the presence of low sequence complexity, intrinsically disordered regions in their structure. These drive clustering/fusion events driven by multivalent interactions between their protein and RNA components, with G3BP1 acting as a key node for promoting RNA-protein, protein-protein and RNA-RNA interactions, ultimately resulting in liquid-liquid phase separation (LLPS) and SG formation (Corbet and Parker, 2019; Wang et al., 2018). SGs are highly dynamic, rapidly assembling to sequester the bulk content of cytoplasmic mRNAs, and dissolving upon stress resolution to release stored mRNAs for future translation (Matheny et al.; Namkoong et al., 2018).

In addition to a general role during the inhibition of translation, SGs can also be stress-selective in composition and function (Aulas et al., 2017). Recent studies have proposed they may adopt non-homogeneous structures with variable compositions dependent on the stress and have proposed classifying SGs into 3 types (Hofmann et al., 2020). Type I canonical SGs form via eIF2α-dependent pathway while type II SGs assemble following eIF2α-independent inhibition of translation. In contrast, type III SGs lack eIFs and are associated with cellular death (Reineke and Neilson, 2019). This suggests that compositionally heterogeneous SGs support specialized functions promoting survival or pro-death outcomes. By sequestering specific proteins, SGs alter the composition and concentration of cytoplasmic proteins, which in turn can change the course of biochemical reactions and signalling cascades in the cytosol (Riggs et al., 2020). Moreover, mutations impacting SG clearance or dysregulating LLPS, can lead to persistent or aberrant SGs, which are increasingly associated with neuropathology, in particular amyotrophic lateral sclerosis (ALS) and related diseases (Wolozin and Ivanov, 2019). Many SG proteins are also aberrantly expressed in tumours and SGs are exploited by cancer cells to adapt to the adverse conditions of the tumour microenvironment (Anderson et al., 2015).

Importantly, SGs are at a crossroads between intracellular signalling, antiviral responses and translation control through concentrating key signalling and cytoplasmic sensors or effectors of innate immunity (Eiermann et al., 2020). Recent work has proposed that SGs exert antiviral activities by providing a platform for antiviral signalling (Eiermann et al., 2020). A well-known antiviral response is the induction of type I interferons (IFNs). During viral replication, double-stranded RNA replication intermediates can be recognized by cytoplasmic sensors such as RIG-I-like receptors (RLRs) or the eIF2α kinase PKR to amplify the IFN response and create an antiviral state (Eiermann et al., 2020; Mateju and Chao, 2021). Multiple IFN signalling molecules, including PKR, MDA5, RIG-I, PKR, and TRAF2, can be recruited to SGs, and this localization has been suggested to regulate their activity (Eiermann et al., 2020; Mateju and Chao, 2021). Furthermore, SGs or specific antiviral SGs (avSG) have been proposed to play a role in antiviral signalling as key signalling proteins including MDA5 and PKR are known to localise to SGs and SG formation is involved in PKR activation (Eiermann et al., 2020; Mateju and Chao, 2021). Because of this proposed role in antiviral signalling and impact on cellular protein synthesis which they rely on, many viruses have evolved strategies to antagonize or exploit SGs, for example by cleaving or repurposing SG-nucleating proteins during infection or impairing the eIF2α sensing pathway (Gaete-Argel et al., 2019). Among these, G3BP1 is a prime target and it is proteolytically cleaved by enterovirus and calicivirus proteases, sequestered by the alphavirus nsp3 protein or repurposed during dengue and vaccinia virus infection (Gaete-Argel et al., 2019). Interestingly, even closely related viruses use different strategies to counteract SGs. Indeed, we previously demonstrated that while Feline Calicivirus (FCV) disrupts the assembly of SGs by inducing G3BP1 cleavage through its 3C-like protease, infection with the related Murine norovirus (MNV) has no impact on G3BP1 integrity (Brocard et al., 2020; Humoud et al., 2016). Instead, viral proteins interact with G3BP1 resulting in its re-localization to replication complexes and in the reshaping of its interactions with cellular partners, repurposing G3BP1 as viral translational enhancer (Hosmillo et al., 2019).

Previous analysis of FCV infection revealed that some uninfected cells near infected cells displayed G3BP1 foci. Herein, we demonstrate that infected cells communicate to nearby bystander cells resulting in the assembly of paracrine granules (PGs). Importantly, while PGs and arsenite-induced SGs share many components such as mRNAs of similar functional families and some resident proteins, PGs exhibit specific features. First, their assembly-disassembly pattern are different, PGs assembly can occur in the absence of the SG scaffold G3BP1 and their disassembly is faster. Second, despite being associated with translational shut-off their assembly is insensitive to cycloheximide and blocking of mRNAs onto polysomes. Finally, PGs induction impairs viral replication suggesting a role in preventing or reducing viral propagation. Therefore, we propose a model in which the assembly of PGs is a pro-survival events resulting from stress signals sent from infected cells to the nearby environment.

## Results

### FCV infection induces granules formation in paracrine manner

We previously reported that FCV infection impairs SGs accumulation in infected Crandell-Rees Feline Kidney (CRFK) cells through NS6-mediated proteolytic cleavage of the SG scaffolding protein G3BP1 (Humoud et al., 2016). Intriguingly, this also revealed that a small fraction of uninfected cells assembles SG-like foci during infection at low multiplicity of infection (Fig. 1A). This suggests a possible paracrine induction of SG-like foci assembly that could contribute to impairing viral replication or propagation in uninfected cells. To test this hypothesis, we generated virus-free supernatant (VFS) from FCV-infected or mock-treated cells (Fig. 1B). Briefly, the cell culture supernatant was collected, and the viral particles were precipitated using PEG3350 and NaCl and removed via ultra-centrifugation and UV-inactivation, as showed in Fig. 1B. The VFS generated was assayed for the presence of infectious viral particles by measuring viral titres after incubation with CRFK cells for up to 72 h using TCID_50_ assays, which confirmed the removal of infectious particles (data not shown). We then tested the ability of the VFS to induce SG assembly in either CRFK or U2OS cells expressing GFP-G3BP1. Cells were stimulated for 1h with VFS or arsenite (ARS), which induces SG by activating the eIF2α kinase HRI (Taniuchi et al., 2016), fixed and labelled with anti-G3BP1 to detect SG assembly. As expected, ARS treatment resulted in the assembly of SGs reflected by the accumulation of G3BP1 into cytoplasmic foci (Fig. 1C). Similarly, treatment of both CRFK and U2OS cells with VFS resulted in the formation of G3BP1 foci in the cytoplasm (Fig. 1C). Detailed analysis in U2OS cells revealed that arsenite and VFS treatments resulted in the formation of G3BP1 foci in 85% and 70% of cells, respectively (Fig. 1D). Interestingly, further analysis also revealed that the average size of foci induced by VFS was significant smaller than the size of arsenite-induced SGs induced by arsenite, and that the number of foci per cell was significantly higher in the VFS-treated cells compared to the arsenite-treated (Fig. 1E), suggesting a different nature of foci induced. We confirmed these results but assessing the presence of another SG resident protein, the RBP FXR1 in SGs. Following stimulation with arsenite or VFS, FXR1 colocalised with G3BP1 in U2OS cells, and analyses further supported that VFS triggered the formation of smaller and more abundant cytoplasmic foci (Fig. 1F and S1 A,B). Thereafter, the VFS-induced G3BP1 foci were therefore named paracrine granules (PGs).

**Figure 1.**
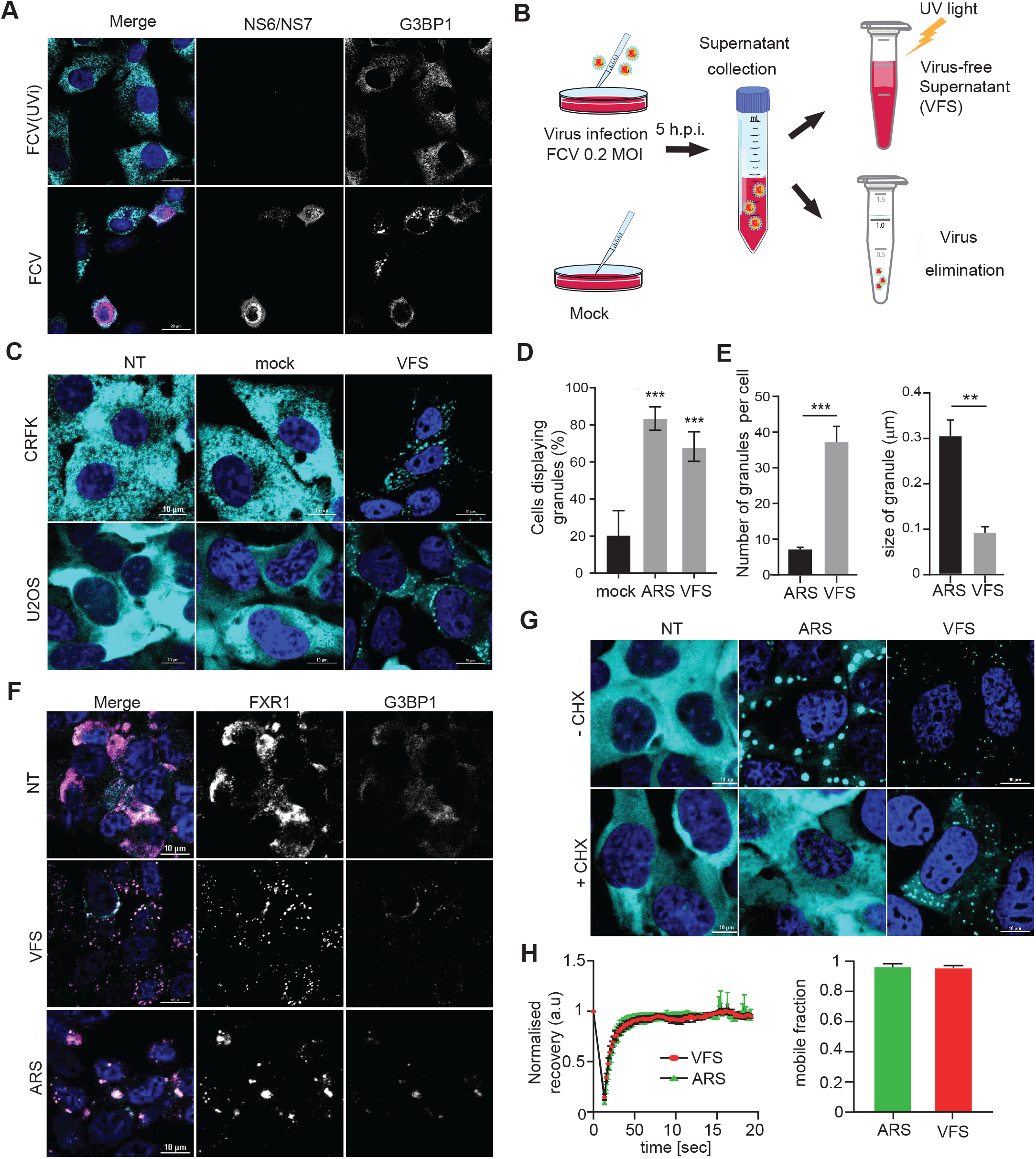
FCV infection results in the formation of paracrine granules. **(A)** Confocal microscopy analysis of CRFK cells infected with FCV (MOI 0.2) or UV-inactivated FCV (FCV(UVi)) for 5h. Samples were stained for the SG marker G3BP1 (cyan) and infected cells detected by immunostaining against FCV NS6/7 (pink) and the nuclei were stained with DAPI (blue). Scale bars, 20μm. **(B)** Schematic representation on virus free supernatant (VFS) preparation procedure. **(C)** CRFK or U2OS GFP-G3BP1 cells were stimulated for 1h with virus free supernatant from a mock or FCV infection and the distribution of G3BP1 analysed by confocal microscopy using detection of endogenous G3BP1 (cyan) in CRFK, or GFP (cyan) in U2OS cells. Non-treated (NT) were used as controls. Nuclei were stained with DAPI (blue). Scale bars, 10μm. **(D)** Representative bar plot (n=3) of the percentage of U2OS cells displaying G3BP1 foci, mean ± SD for 100 G3BP1-positive cells analysed across at least 10 acquisitions. Statistical significance shown above the bars, ***, *P* < 0.001. **(E)** Representative bar plot (n=3) of the number of G3BP1 granules per cells displaying G3BP1 foci and the average granule size, mean ± SD for 100 G3BP1-positive cells analysed across at least 10 acquisitions. Statistical significance shown above the bars, ***, *P* < 0.001, **, *P* < 0.01. **(F)** U2OS GFP-G3BP1 cells were stimulated for 1h with VFS or 0.5mM sodium arsenite (ARS) prior to fixation and the formation of stress granules analysed by confocal microscopy using G3BP1 or FXR1 as markers. Non-treated (NT) were used as controls. Nuclei were stained with DAPI (blue). Scale bars, 10μm. **(G)** VFS-induced G3BP1 aggregates are CHX insensitive. U2OS GFP-G3BP1 cells were treated with ARS or VFS. For forced SG disassembly, cells were treated with 10μg/ml of CHX for 30min (+CHX). The presence of G3BP1 granules was assessed as in **(D). (H)** Fluorescence recovery after photobleaching (FRAP) analysis of GFP-G3BP1 granules in U2OS GFP-G3BP1 cells treated with 0.5 mM ARS or VFS. Plot shows recovery curves as an average of 9 granules for VFS (red) and 7 granules for ARS (green) ± SEM. Images were taken every 3s during recovery. Mean intensity was determined at each time point using ImageJ (It). Mean intensity of GFP within the cell from a different part of the cytoplasm was measured (IBt) to correct for bleaching during image acquisition. Corrected mean intensity at each time point was determined by taking the ratio: It/IBt and used to calculate the diffusion mobility coefficient of GFP-G3BP1.

To further characterize the features of PGs, their susceptibility to cycloheximide (CHX) treatment was determined. The assembly of canonical SG is dependent on the shuttling of mRNAs dissociating from polysomes into SGs. By binding to ribosomes and preventing mRNAs release, CHX inhibits SG formation (Kedersha et al., 2000). U2OS GFP-G3BP1 cells were pre-treated with CHX and then treated with either arsenite or VFS. As expected, CHX impaired the assembly of canonical SGs, however it was unable to block PG formation following VFS treatment (Fig. 1G), strongly suggesting that PGs form in response to different stimuli than canonical SGs.

We next examined the kinetic of assembly-disassembling of PGs compared to SGs using time-lapse confocal microscopy (Fig. S1). First, G3BP1 foci formation was recorded by collecting images every 10 minutes for 180 minutes, and the disassembly of these foci upon stressor removal recorded in a similar manner. As showed in Fig. S1C, PGs formed faster than the SGs, with 10 minutes stimulation with VFS sufficient to induce PGs assembly, whereas canonical SGs required a longer time, between 30 to 40 minutes of arsenite treatment, in order to form completely. More strikingly, following foci disassembly in recovery assay revealed that while canonical SGs required up to 3h to disassemble, PGs completely disappeared within 20 minutes of the t0 stress removal (Fig. S1D), potential reflecting a more dynamic nature of these condensates. To test this further, the internal mobility of GFP-G3BP1 was measured using Fluorescence recovery after photobleaching (FRAP), cells were treated either with VFS or arsenite. Surprisingly, we could not measure any differences in the recovery of GFP-G3BP1 fluorescence after photobleaching, and thus in G3BP1 mobility between the two conditions (Fig. 1H). Overall these data suggest that VFS from infected cells induce the formation of PGs that display specific features of assembly and disassembly dynamic, different to canonical SGs.

### Proteomics analysis of isolated PG reveals a distinct composition from SGs

Previous studies have suggested in response to different stresses, SGs recruit specific resident proteins and that this SG heterogeneity may be important for their cellular functions (Reineke and Neilson, 2019; Youn et al., 2019). To explore this, we took advantage an affinity-based SG isolation procedure we recently developed to characterise their proteome (Jain et al., 2016). To this end, U2OS GFP-G3BP1 cells were treated with VFS for 1h to induce SG assembly. G3BP1 interactors and SG cores were then enriched by sequential centrifugation to generate a granule enriched fraction (GEF) and purified by immunoprecipitation using antibodies to GFP (to trap GFP-G3BP1) or IgG (as a control) followed by pull down with Protein A-conjugated Dynabeads as previously described and summarised in Fig. 2A (Brocard et al., 2020; Jain et al., 2016). Epifluorescence microscopy analysis then confirmed the isolation on anti-GFP beads of SG cores, while no GFP signal could be detected in the control IgG immunoprecipitation (Fig. 2B). To characterize the identity of PGs resident proteins, mass spectrometric analysis by LC-MS/MS was performed on proteins eluted from the beads and analysed using MaxQuant. 667 proteins were detected with a false-discovery rate (FDR) of less than 5%, 191 proteins displayed at least two different peptides and applying a filtering criterion of ≥1 Log_2_-fold changes of immunoprecipitated proteins compared to the respective control IgG conditions, we finally identified 110 proteins enriched in the PGs (Table S1).

**Fig 2.**
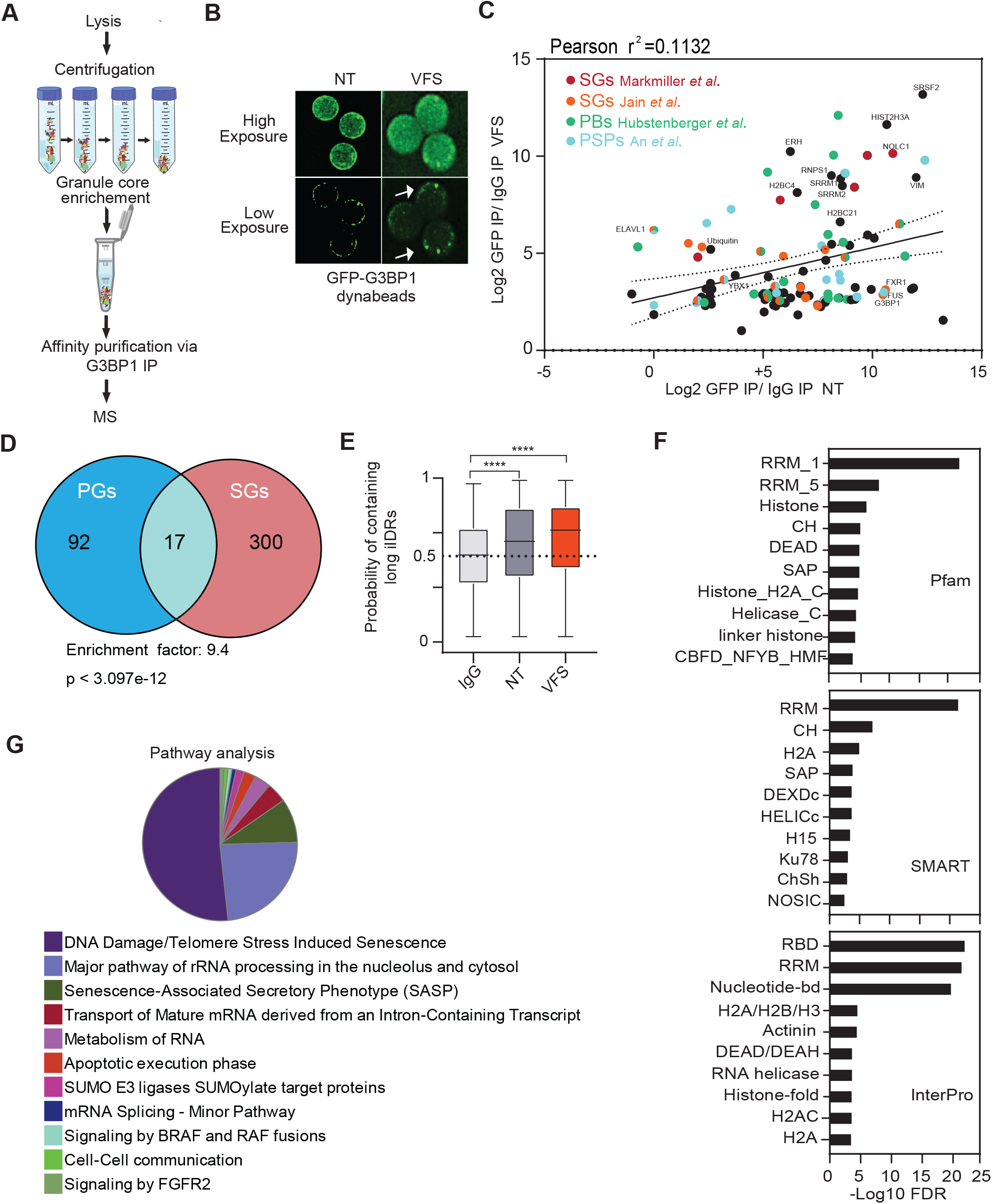
Proteomic analysis reveals differences between VFS-induced PGs and ARS-induced SGs. **(A)** Schematic representation of the granule isolation procedure. **(B)** Dynabeads bound to GFP analysed by epifluoresence microscopy in ARS or VFS treated U2OS GFP-G3BP1 cells; bead-bound G3BP1 granules are indicated by white arrows. High-and low-level exposure are shown. **(C)** Scatterplot of 110 proteins enriched in the PGs over the control (Log2 transformed for each IP ratio relative to control IgG). Proteins previously identified as components of other biocondensates (SGs, P-bodies: PBs, Paraspeckles: PSPs, Membraneless organelles: MLOs) are indicated in colour. **(D)** Venn diagram between 110 proteins identified in PGs and 317 in SGs (from Jain *et al* 2016). The representation factor showed the enrichment over the expected and p-value is based on the cumulative distribution function (CDF) of the hypergeometric distribution of the data set over the mouse proteome. **(E)** Box-plots of the average score of the probability of a protein containing a long intrinsically disordered region determined using SLIDER in the GFP versus IgG IPs for VFS-treated or control (NT) cells. Statistical significance shown above the bars, ****, *P* < 0.0001. **(F)** Bar plot of the most enriched protein domains in the PGs discovery analysis using Pfam, SMART and InterPro. **(G)** GO pathway analysis of the 110 resident proteins isolated in the PGs using cytoscape.

We next compared the PGs protein composition with that of previously published ARS-induced SGs (Jain et al., 2016; Markmiller et al., 2018) or other biocondensates such as P-bodies (PBs) (Hubstenberger et al., 2017) and paraspeckles (PSPs) (An et al., 2019) (Fig. 2C, Table S2). This highlighted that the composition of PGs is widely different from these other biocondensates. More specific comparison of the previously established arsenite-induced SG proteome in U2OS cells revealed PGs share a number of components with canonical SGs, as showed in Fig. 2D, Table S3 (Jain et al., 2016), with 17 out of 110 PG proteins displaying a significant enrichment in SGs (946 fold p-value: p < 3.097e-12). These include RBPs such as FXR1 and ELAVL1, and RNA helicases such as DDX1 and DDX21. We further confirmed the cellular distribution of some of these targets using immunofluorescence following treatment with either ARS or VFS. The known SG residents ELAVL1, FXR1 or UBAP2L colocalised with G3BP1 both in PGs and SGs (Fig. S2). In contrast, hnRNPK, THRAP3 and RBMX colocalised with G3BP1 in PGs but not in arsenite-induced SGs, confirming the assembly of compositionally distinct foci (Fig. S3).

Recent studies have proposed a model for SG formation structured around the interplay of a large interaction network of mRNAs and key nucleator proteins enriched in intrinsically disordered regions (IDRs), Prion-like domains (PrLDs) and RNA-binding domains (RBDs) (Guillen-Boixet et al., 2020; Sanders et al., 2020; Yang et al., 2020). We used the online tool SLIDER, which predicts whether a protein sequence has long disordered segments (at least 30 consecutive disordered residue) to analyse to screen the PG proteins (Peng et al., 2014). We observed an increased in proteins with long IDRs in the G3BP1 IP compared to the IgG pulldown (Fig. 2E). Moreover, using STRING for interaction analysis we performed *in silico* screening in order to identify which protein domains were most enriched in the PGs (Szklarczyk et al., 2019). RBDs and RNA recognition motifs (RRM) were the most enriched domains within the reported top 10 protein domains identified across three different databases Pfam, SMART and InterPro confirming in (Fig. 2F). Finally, to better characterise the common feature of the 110 PG proteins, we performed a Gene Ontology (GO) enrichment analysis with Cytoscape using GlueGo and the online platform “Metascape” (Zhou et al., 2019). This confirmed that the overrepresentation in the PG composition of RNA binding proteins (RBPs), for example factors involved in RNA splicing (GO:0008380-GO Biological Processes), RNA localization (GO:0006403), mRNA catabolic process (GO:0006402), and it highlights the composition in focal adhesion (GO:0005925 GO Cellular Components) and ribonucleoprotein granule (GO:0035770). A further GO screening was carried out identify possible pathways (KEGG and Reactome) activated by VFS stimulation (Fig. 2G). Interestingly, metabolism of RNA (R-HSA:8953854) and cell to cell communication (R-HSA:1500931) were identified among the predicted pathways, fitting with a paracrine mechanism of induction. We also identified pathways induced by stress like senescence and BRAF signalling (R-HSA:6802952) or other paracrine pathway like FGFRs (R-HSA:5654738) (Table S4).

### PGs assembly results in the condensation of functional classes of mRNAs similar to those found in SGs

Previous studies have underpinned the roles of mRNA species in the assembly of SGs, showing they support the network of weak and transient interactions required during condensation (Van Treeck et al., 2018). In addition, RNAs can condense on their own resulting in SG-like condensate, with almost identical mRNA contents, in the absence of protein (Van Treeck et al., 2018). Moreover, while SGs are thought to sequester a wide variety of cytoplasmic mRNAs, specific transcripts excluded from sequestration can drive a specific translational programme to adapt to stress (Khong et al., 2017; Namkoong et al., 2018). Thus, to further characterise the PG/SG differences, we analysed the RNA contents of PGs using RNA-seq. To this end, we used the granule-enriched fraction (GEF) given as it has previously been shown to accurately reflect the RNA content of isolated SGs (Namkoong et al., 2018). As outlined in Fig. 3A, U2OS GFP-G3BP1 cells were treated with VFS, or untreated, lysed and following preparation of GEF fraction, RNAs were then purified, sequenced by Illumina sequencing (PG transcriptome), and compared to the total RNA (total transcriptome).

**Fig 3.**
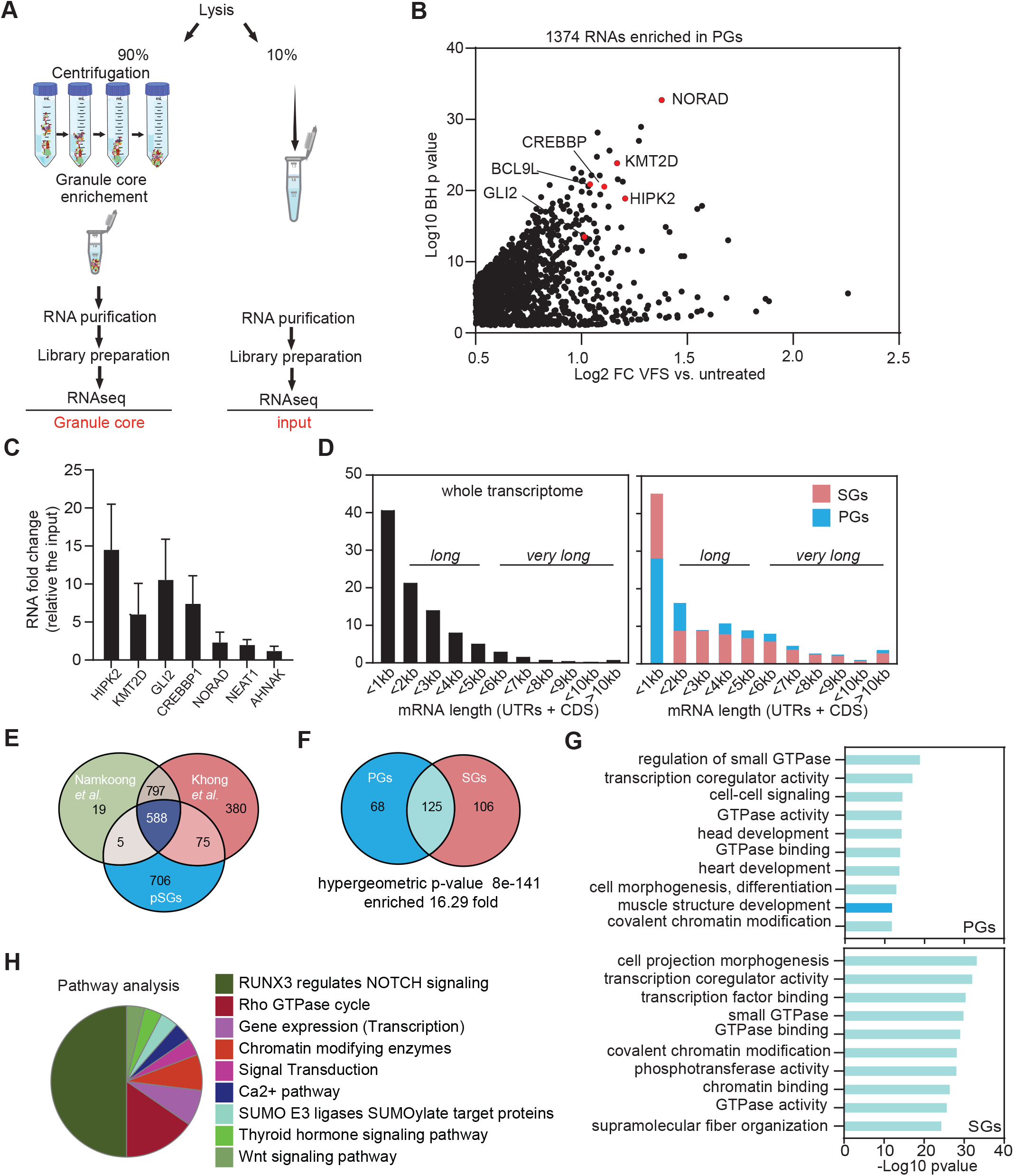
Transcriptomic analysis reveals differences between VFS-induced pGs and ARS-induced SGs. **(A)** Schematic representation of the procedure for total and PG transcriptome analysis in VFS or untreated cells. **(B)** Volcano plot showing statistically significantly enriched differentially expressed RNAs (Log2 fold change of VFS versus non-treated) in the PGs. **(C)** Transcript levels of PG resident mRNAs were quantified via RT-qPCR relative to untreated and normalized to the individual total level of each RNA. Error bars represent SEM (n = 3) and statistical significance shown above the bars, * P < 0.05, *** P < 0.005. **(D)** Comparison of the mRNA length for transcripts enriched in PGs and arsenite-induced SGs (from (Khong et al., 2017)) compared to total transcriptome distribution. **(E)**. Venn diagram comparison of mRNAs enriched in PGs and arsenite-induced SGs (from (Khong et al., 2017 and Namkoong, 2018 #21)). **(F)**. Venn diagram comparison of the GO terms (molecular function and biological process) enriched in the mRNAs enriched in pGs or SGs upon VFS-stimulation or sodium arsenite (ARS) treatment (from (Khong et al., 2017)). **(G)** Comparison of the top 10 GO terms enrichment (molecular function and biological process) for mRNAs enriched in PGs (dark blue) or SGs (pink). GO terms overlapping both conditions are in light blue. **(H)**. Clustering by functional pathways, identified from gene ontology analysis (KEGG/Reactome) of the mRNAs enriched in VFS-treated versus non-treated U2OS cells.

Total transcriptome analysis identified 15698 transcripts overall and 7450 were selected for further analysis with a significant BH p-value <0.05 (Table S5). We analysed the coverage of the different RNA species present in the RNA-seq data set and we did not observe any rRNA accumulation confirming that our sample were efficiently rRNA-depleted, and as expected the majority of the RNA species identified, 95.9%, were protein encoding, with long non-coding RNAs (lncRNAs) representing 2.5% and the remaining enriched for other non-processed RNAs, small nucleolar RNAs (snoRNAs), small nuclear RNAs (snRNAs) and microRNAs (miRNAs) (Fig. S4A). We further looked at different features of RNA, like length and GC content. We observed that 41% of mRNAs were less than 1kb-long and the majority of RNAs (67.3%) contained less the 50% GC content (Fig. S4B). Furthermore, we performed a differentially expression analysis comparing the RNAs expressed (transcriptome) in the VFS versus untreated cells used as control, we identified 712 RNAs that were up regulated to be at least and above 0.5 Log_2_ CPM of RNAs in VFS-treated relative to the untreated cells and 2510 RNAs were down-regulated (Fig. S4C, Table S6). This analysis revealed that the majority of the RNAs were downregulated in VFS versus the control, including many lncRNAs such as NORAD and NEAT1(Fig. S4D). Both up and down-regulated microRNAs could be identified, while the majority of lncRNAs were downregulated. We validated some of the transcript RNAs via RT-qPCR analysis (Fig. S4E), confirming the inhibition of gene expression induced by VFS treatment, and observed a strong downregulation of BCL9L, HIPK2, CREBBP1, GLI2, NORAD and NEAT1. Interestingly, the majority of mRNAs that were not downregulated encoded for ribosomal proteins previously shown to exhibit a stable/long half-life such as RPS18, RPL9 (Fig. S4E).

Functional GO analysis (molecular function and biological process) revealed that the majority of upregulated RNAs are involved in the oxidative phosphorylation (GO:0006119), such as ATP synthase, cytochrome c oxidase and NADH-ubiquinone oxidoreductase; mitochondrial (GO:0006839) and translation-like ribosome components (hsa03010) (Fig. S5A). We further compared GO terms enrichment for the top 500 significant RNAs identified in VFS or arsenite-treated U2OS cells and U2OS G3BP1-GFP cells (Fig. S5B,C, Table S7) (Khong et al., 2018). Analysis of the most enriched GO terms revealed a small overlap, suggesting that the overall pathways activated by these two types of stresses are different. The comparison of the top 10 summary GO terms enrichment showed that the VFS induces an effect mainly at transcription regulation level (GO:0003712, GO:0008134, GO:0001227) in contrast to mRNA metabolism for ARS (GO:0022613, GO:0006402, GO:0031145).

In contrast, SG transcriptome analysis identified 1374 transcripts with a 0.5 Log_2_ fold change above the background (untreated cells) (Fig. 3B and Table S8). As expected most of the RNAs identified encoded for protein (92.3%), where the remaining transcripts were identified to be 3.7% lncRNAs, 1.5% microRNAs and the rest for RNAs like snoRNA, and process and unprocessed pseudogenes. We confirmed by RT-qPCR some of the most enriched RNAs identified in PGs including HIPK2, CREBBP1, GLI2 and KMT2D, whereas lncRNAs as NORAD, NEAT1 and AHNAK were only slightly enriched (Fig. 3C).

We next analysed whether enriched transcripts presented common features, such as length or GC contents, that could indicate their ability to form large RNPs. We observed that only 28.5% of PGs mRNAs in PGs were “short” or less than 1 kb, with the majority of mRNAs enriched, 46.7%, “long” or between 1 and 5 kb, and 24.8% was “very long” or above 5 kb. This was surprisingly different compared with the mRNAs enriched in canonical SGs where the majority of the mRNAs were “short” 45.7%, only 33.9% were “long” and just 20.4% were “very long” mRNAs (Khong et al., 2018) (Fig. 3D). The majority of RNAs enriched in PGs, 91.2%, displayed less than 50% GC content, reflecting not very structured RNAs.

Next, comparison of RNAs enriched in arsenite-induced SGs and PGs revealed only 50.7% overlap between the two datasets (Fig. 3E). Therefore, we compared the GO terms enrichment for both SGs and PGs RNAs and for the SGs induced by arsenite in U2OS GFP-G3BP1 cells (Fig. 3F). This revealed a significant overlap between the two classes of condensates (16.3 fold), suggesting that although the 50% of RNAs enriched in the PGs were different from the RNAs enriched in the SGs, they shared significant functional roles (Fig. 3F,G, Table S9). The top 10 GO terms enriched in PGs involved the small GTPase mediated signal transduction (GO:0007264) and cell junction organization (GO:0034330) were also significantly enriched in the 125 GO terms in common between PGs and SGs, and *vice versa* (Fig. 3G, Table S10). Finally, analysing the most enriched pathways (Reactome and KEGG), using the filter at p-value<0.01, highlighted that the cell-cell communication pathway (RUNX3 -NOTCH) represented 50% of the enriched GO terms (Fig. 3H, Table S11). Overall these suggest that VFS treatment induce global changes in the U2OS transcriptome that are different from those observed during arsenite-induced oxidative stress, but that similar functional classes of mRNAs are enriched in SGs and PGs.

### N6-methyladenosine modified RNAs and ARE motif are enriched in PGs

To investigate whether specific RNA motifs were enriched in the PG transcriptome we selected the top 500 most significantly enriched mRNAs and analyse motifs enrichment using the MEME and DREME (Bailey et al., 2009). The top five motifs enriched in the PGs are displayed in Fig. 4A, with corresponding “E-value” for each identified motifs and included GA-GA[U/G/A]GA, CGC[G/C/A]GC[G/C]G, UAUUU[U/A]U[U/A], [G/C]AGCAGC[U/A] and U[G/A]UAUAU[G/A]. We then correlated these with RBPs binding using the MEME-built in tomtom tool built to identify putative RBPs previously described to bind each specific given motif (Fig. 4A). Interestingly this revealed that some of the RBPs identified in as resident PG proteins by our proteomic analysis are targets of the enriched RNA motifs, such as ELAVL1, SRSF2 and SRSF7 and G3BP2 which shares similar binding motif as G3BP1 (Edupuganti et al., 2017).

**Figure 4.**
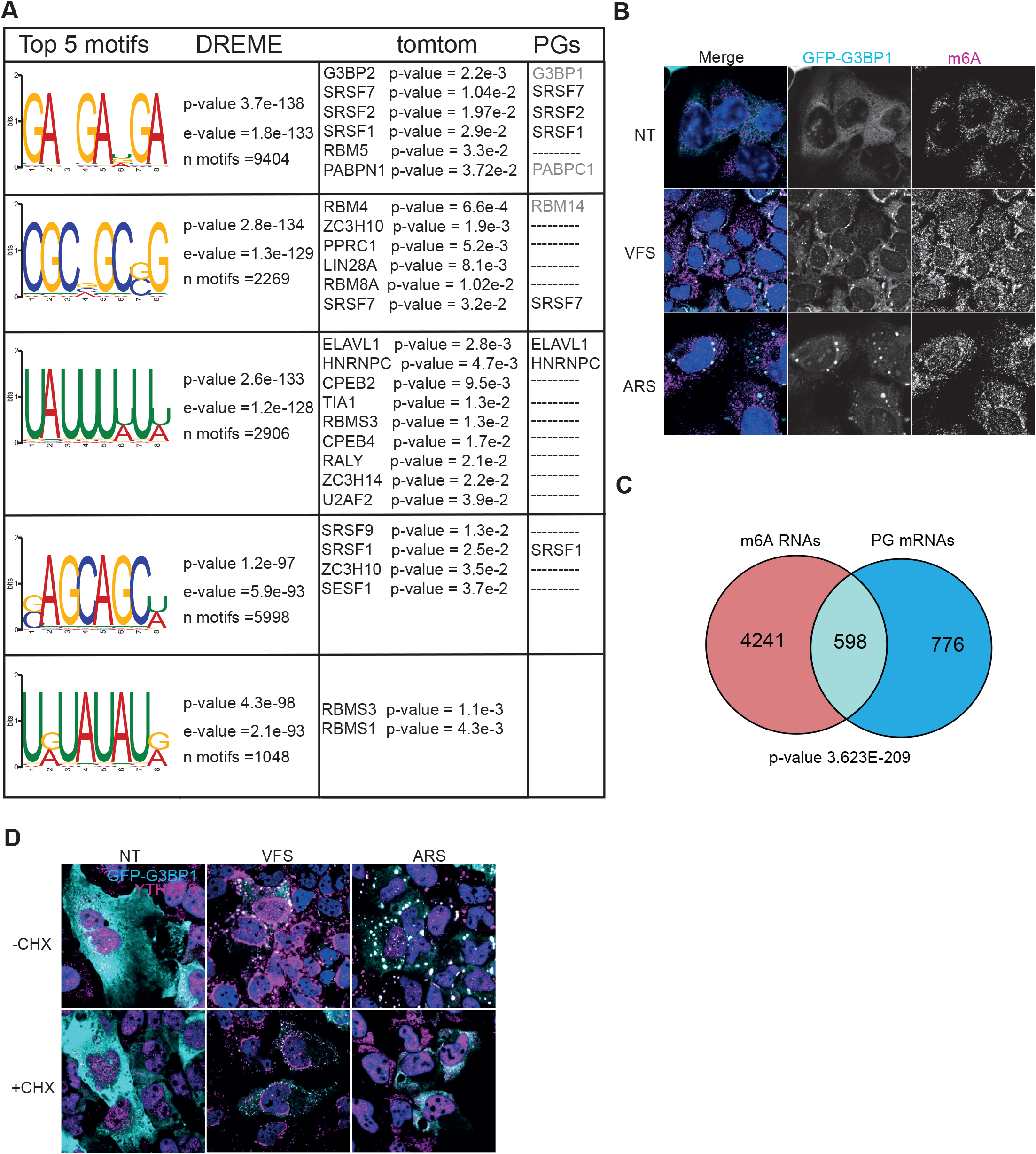
N6-methyladenosine modified RNAs are enriched in PGs. **(A)** Top 5 RNA motifs identified from DREME analysis of the PG transcriptome, with corresponding putative RBP targets identified from tomtom analysis of these motifs and the corresponding proteins identified by proteomic as pG components. **(B)** U2OS GFP-G3BP1 cells were stimulated for 1h with VFS or 0.5mM sodium arsenite (ARS) prior to fixation and the formation of PGs/SGs analysed by confocal microscopy. Non-treated (NT) were used as controls. Cells were stained with GFP (cyan) as PG marker and m6A (magenta) as N6-methyladenosine modified RNAs marker. Nuclei were stained with DAPI (blue). Scale bars, 10μm. **(C)** Venn diagram between 1374 RNAs identified in PGs and 4839 m6A-edited mRNAs localised to SGs (from Anders *et al* 2018). The representation factor showed the enrichment over the expected and p-value is based on the cumulative distribution function (CDF) of the hypergeometric distribution of the data set over the human genome. **(D)** The presence of G3BP1 granules was assessed as in **(B)** following forced SG disassembly using 10μg/ml of CHX for 30min (+CHX). Cells were stained with GFP (cyan) and YTHDF3 (magenta). Nuclei were stained with DAPI (blue). Scale bars, 10μm.

Additionally, it has been proposed N6-methyladenosine (m6A) RNA modifications play a role in triaging mRNAs from the translatable pool into SGs (Anders et al., 2018). Upon binding of these target mRNAs, their m6A reader proteins, the YTH domain family proteins (YTHDFs), undergo LLPS and promote SG formation (Elbaum-Garfinkle, 2019; Yang et al., 2020). This overlap points to a possible link between m6A modification, translation under stress conditions and possible recruitment to SGs (Zhou et al., 2015). To determine if m6A-modified RNAs were enriched in PG, we performed immunofluorescence in U2OS GFP-G3BP1 cells treated either with VFS or arsenite and analysed the cellular distribution of m6A mRNAs using a specific antibody previously used to pull-down and identify these mRNAs (Anders et al., 2018). As shown on Fig. 4B, both ARS and VFS treatment resulted in colocalization of m6A signal with G3BP1. Next, we compared the list of m6A-modified mRNAs localised to SGs following arsenite treatment from Anders *et al* (Anders et al., 2018) with PG resident mRNAs (Fig. 4C). We observed a 3.8 fold enrichment between the two groups, with a significant overlap (hypergeometric p-value = 3.6e-209), confirming that a significant proportion of PGs mRNAs could be m6A-modified. Since m6A-mRNA triage is thought to be driven by YTHDF3, localising to SGs during stress, we performed immunofluorescence to investigate its cellular localisation following VFS treatment. This confirmed that YTHDF3 colocalised with G3BP1 in the PGs, and that unlike in arsenite-induced SGs this localisation is unsensitive to CHX treatment (Fig. 4D). We also notice that YTHDF3 displayed a peculiar distribution around the nucleus not observed with arsenite-induced stress. Overall, these data suggest that like SGs, PGs may condensate m6A-modified mRNAs and their reader protein YTHDF3

### G3BP1 is not essential for PGs assembly

With strong indication that PGs and SGs share common features but exhibit fundamental differences, we next investigated the importance of G3BP1 for PGs assembly. G3BP1 is central and acts as a molecular switch for SG assembly (Yang et al., 2020). Following translation inhibition, the increase in cytoplasmic mRNAs facilitate the clustering of G3BP1 through protein-RNA interactions into networked condensates. These condensates recruit additional client proteins that promote LLPS and SG maturation. Importantly, during SG nucleation cores can also assembly around the RBP UBAP2L, even in absence of G3BP1 (Cirillo et al., 2020). First, we confirmed the presence of UBAP2L in FXR1 and G3BP1 positive PGs (Fig. 5A). To investigate G3BP1 requirement for PG assembly, U2OS WT or G3BP1 knockout (ΔΔ G3BP1) cells were treated with VFS or arsenite and stained for the PG/SG core markers UBAP2L and FXR1. In untreated WT or ΔΔ G3BP1 cells, both FXR1 and UBAP2L remained diffused in the cytoplasm (Fig. 5B). In contrast VFS treatment resulted in the assembly of PGs that contained both FXR1 and UBAP2L in either U2OS WT or ΔΔ G3BP1 cells (Fig. 5B). These results further highlight the heterogenous nature of PGs whose assembly, unlike SGs, is independent from the presence of G3BP1. Previously, small SG-like punctate known as RNase L-bodies (RLB) were shown to assemble independently from G3BP1 in response to non-self sensing by RNase L (Burke et al., 2020). To ascertain whether PGs and RLBs are in fact similar aggregates we compared their proteins composition. Despite a higher proportion of shared components, and a higher enrichment factor, with RLBs rather than SGs or other biocondensates, this revealed a clear distinct composition of PGs and RLBs, suggesting these are distinct bodies (Fig. S6 and Fig. 2). RLBs and PGs are also distinct in their timing of assembly since we observed PGs form rapidly within 10-20 min of VFS exposure, while RLBs form over 1-2 hours and are preceded by the formation of SGs that then remodel into RLBs (Burke et al., 2020).

**Figure 5.**
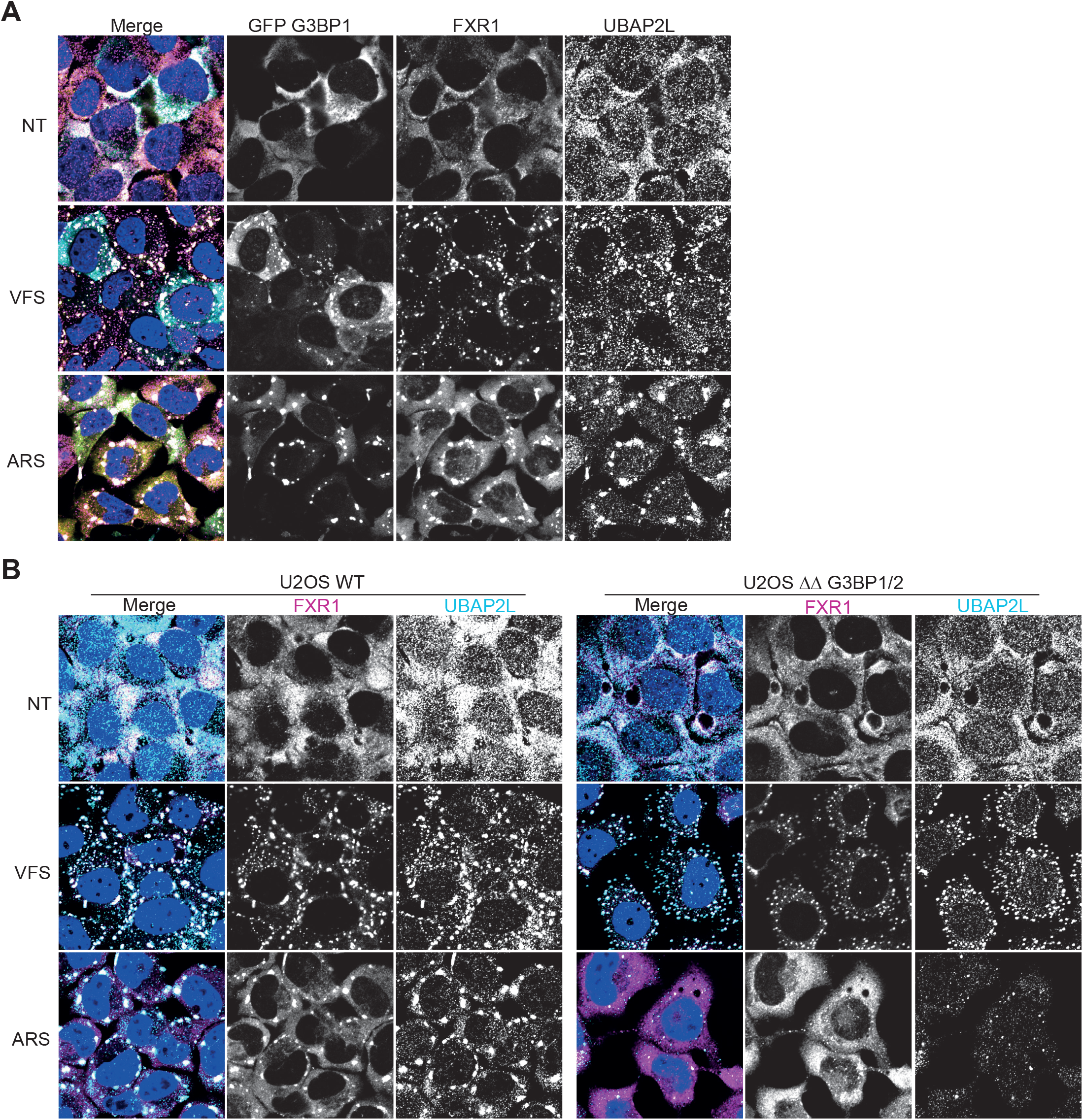
G3BP1 is not an essential for PG assembly. **(A)** U2OS GFP-G3BP1 cells were stimulated for 1h with VFS or 0.5mM sodium arsenite (ARS) prior to fixation and the formation of PGs/SGs analysed by confocal microscopy. Non-treated (NT) were used as controls. Cells were stained with GFP (cyan), FXR1 (magenta) and UBAP2L (gold) as PG/SG markers. Nuclei were stained with DAPI (blue). Scale bars, 10μm. **(B)** Wild-type (WT) or ΔΔ G3BP1 U2OS cells were treated as in **(A)** and stained FXR1 (cyan)) and UBAP2L (magenta) as PG/SG markers. Nuclei were stained with DAPI (blue). Scale bars, 10μm.

### PGs assembly is coupled to translational stalling and activation of stress signalling pathways

Canonical SGs are assembled in response in response to translational stalling that can be triggered by eIF2α-dependent or independent pathways and result in the storage of the bulk of cytoplasmic mRNAs until stress is resolved. Thus, we investigated whether PGs assembly is also coupled to translational inhibition. To this end, we determined global translational efficiency using single cell analysis by measuring the incorporation of puromycin, a tRNA structural mimic which specifically labels actively translating nascent polypeptides and causes their release from ribosomes. Puromycylated native peptide chains are then detected with anti-puromycin antibodies and confocal microscopy (Fig. 6A). As expected, quantification of the puromycin signal intensity revealed a strong decrease in protein synthesis following treatment with arsenite in U2OS cells (Fig. 6A,B). Treatment of U2OS cells with VFS also resulted in impaired puromycin incorporation and inhibition of protein synthesis (Fig. 6B). Therefore, PGs assembly, like SGs, is coupled to translational shut-off. To understand the cellular signalling pathways responsible for communicating this stress we then analysed the activation of several stress-associated signalling proteins using immunoblotting (Fig. 6C). As expected, ARS treatment resulted in the phosphorylation of eIF2α, which occurs through the kinase HRI. VFS treatment also induced eIF2α phosphorylation, relative to the total level of total eIF2α protein level (Fig. 6C). None of the hallmarks of eIF2α-independent signalling activation via mTOR could be detected, with no phosphorylation of AKT, S6K and mTOR but there was not a clear phosphorylation inhibition observed. In contrast, analysis of MAPK signalling pathway revealed that VFS treatment resulted in a strong increase in ERK phosphorylation, but not p38 or eIF4E. These results suggest that PGs assembly is coupled to translational inhibition and activation of stress-related signalling pathways such as eIF2α and ERK1/2.

**Figure 6.**
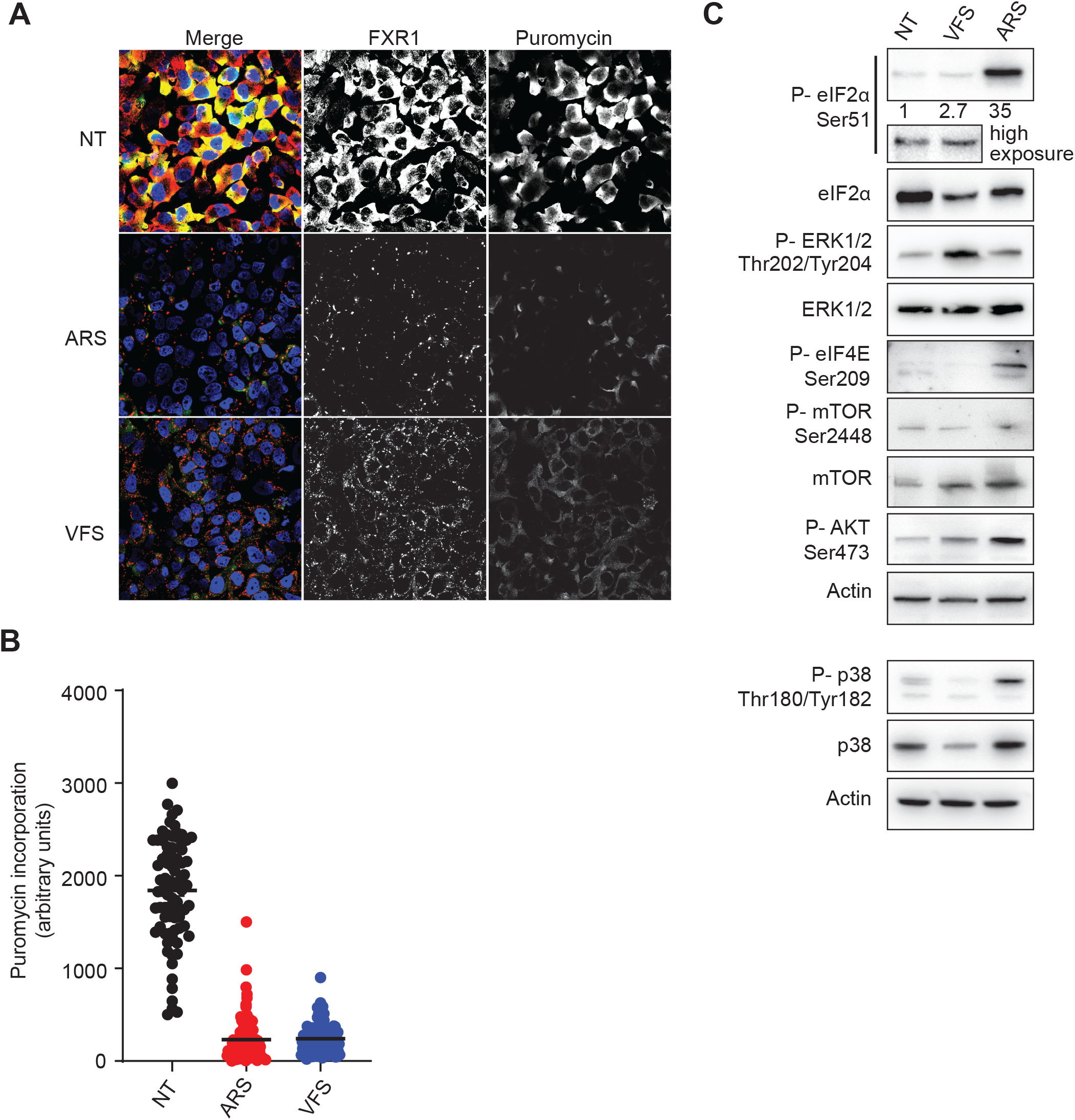
PG assembly is associated with global shutoff in translation and activation of intracellular signalling pathways. **(A)** U2OS cells stimulated for 1h with VFS or 0.5 mM sodium arsenite (ARS) were incubated with 10μg/ml of puromycin to label the nascent polypeptidic chains prior to fixation. Non-treated (NT) cells were used as a negative control. Puromycin-labelled chains were visualised by immunostaining against puromycin (green) and PG/SGs cells were detected by immunostaining against FXR1 (red). Nuclei were stained with DAPI. Scale bars, 10μm. **(B)** Representative scatter plots of *de novo* protein synthesis measured by fluorescence intensity of the puromycin signal (n=3), a.u. arbitrary units. **(C)** Representative Western Blot analysis (n=3) of stimulated as in **(A)**. Specific antibodies used are indicated to the left.

### PGs playing an antiviral response

The formation of PGs was first observed during infection of CRFK cells with FCV in bystander cells. Recent work has proposed that SGs exert antiviral activities by providing a platform for antiviral signalling (Eiermann et al., 2020). Multiple IFN signalling molecules, including PKR, MDA5, RIG-I, PKR, and TRAF2, can be recruited to SGs, and this localization has been suggested to regulate their activity (Eiermann et al., 2020). Therefore, we hypothesised that PGs assembly could be part of paracrine signalling to reduce or prevent the spread of FCV to uninfected bystander cells. To test this hypothesis, CRFK cells were treated with increasing amount of VFS, then infected with FCV for 6h at MOI 2, and viral titre measured using TCID_50_ assays. FCV replication was impaired by treatment of CRFK with neat VFS, with 95% reduction in infectious titres, while dilution of VFS rescued viral replication (Fig. 7A). Finally, we tested whether this impairment of viral replication could be explained by the triggering of antiviral factors. Following 1 or 6 h treatment with VFS, total RNA was extracted from CRFK cells and the levels of IFN-α/β, TNFα and IL10 measured by qPCR. Compared to untreated control, VFS did not induce IFN-α/β, TNFα or IL10 expression levels (Fig. 7B). These suggests that the VFS-induced reduction in viral replication may be due to translational stalling rather than activation of antiviral signalling in cells assembling PGs.

**Figure 7.**
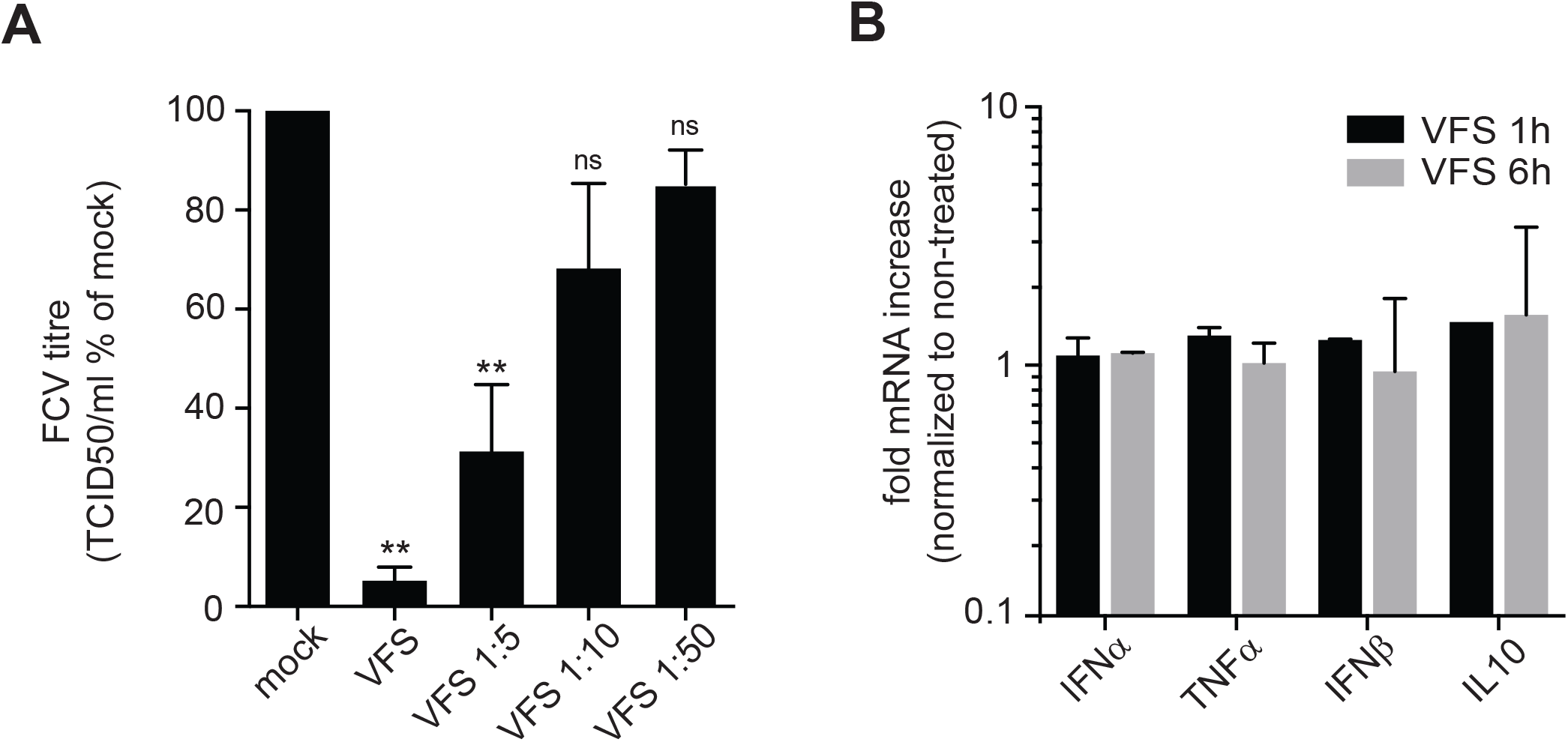
VFS treatment impairs FCV replication in CRFK cells. **(A)** CRFK cells were stimulated for 1h with decreasing amount of VFS and infected with FCV at an MOI of 1. The cells were incubated for 12 h, and the viral titer was estimated by a TCID50 assay. Three separate experiments were analysed by standard t test (**, P<0.01). **(B)** Transcript levels of indicated mRNAs were quantified via RT-qPCR in CRFK cells following stimulation with VFS for 1 or 6h and normalized to non-treated cells. Error bars represent SEM (n = 3).

## Discussion

In the present study we have identify a new type of RNA granules induced in bystander cells near infected cells during FCV infection. Our results support a model in which paracrine signalling during infection results in the activation of stress-related intracellular signalling, including ERK and eIF2α pathways and a shut-off of translation. This is coupled to the assembly of SG-like condensate we named PG. These PGs share some characteristics with the previously identified SGs, PBs, RLBs, and PSPs, but exhibit their own specific nature. We mainly compared the PGs to the canonical SGs induced by arsenite in U2OS cells and observed numerous differences in granule size, number per cell and dynamics. Resident proteins are, like for SGs, enriched in RNA-binding proteins and proteins with disordered domains, yet they condensate distinct functional classes of proteins when compared to SGs. We also characterised the RNA content of these granules. In contrast to SGs, RNAs found in PGs are significantly longer and the identity of concentrated RNAs is different, yet similar functional classes of mRNAs are enriched within SG and PG. Interestingly, we found that these PG-resident RNAs were enriched for RNA motifs recognised by RNA-binding proteins we identified in PGs, such as ELAVL1, SRSF2 and 7 and G3BP2. This raises the possibility that these proteins may play important role during the PG condensation process by driving RNA-protein interactions.

Our analysis of PG assembly/disassembly dynamic revealed that although their assembly followed a similar pattern to that of arsenite-induced SGs, in contrast they disassembled more rapidly with complete clearance within 10 minutes (Fig. S1). This behaviour is reminiscent of the previously characterised cold-shock SGs (Hofmann et al., 2012). In mammalian cells, following cold shock exposure to hypothermia, the PERK-mediated phosphorylation of eIF2α and inhibition of mTOR signalling impairs the initiation of translation and the assembly of preinitiation complexes, resulting in protein synthesis slow-down and triggering SGs assembly (Hofmann et al., 2012; Roobol et al., 2009). While cold shock-induced SGs form over the course of several hours, they disassemble within minutes when cells are returned to ideal growth temperatures. Interestingly, cold shock-induced SGs condense poly(A) mRNA and several known SG resident proteins such as eIF3B, eIF4G, eIF2α, G3BP, PABP, HuR, and TIA1, confirming that these are *bona fide* SGs, however they are smaller in sizes and more abundant than arsenite-induced SGs. Reacting to the reduction in energy associated with hypothermia, AMP-activated protein kinase (AMPK) signalling is further activated to orchestrate cellular adaptation to these stressful conditions. This activation is essential for cold shock-induced SG assembly as pharmacological inhibition of AMPK signalling prevents the assembly of cold shock-induced SGs and the survival of mammalian cells at low temperatures. In own study of intracellular signalling rewiring during PGs assembly did not reveal clear activation of AMPK signalling targets, thus suggesting that PGs and cold shock-induced SGs are associated with distinct pathways of cellular adaptation to stress (Fig. 6D).

In addition to PKR activation previously mentioned, the OAS/RNase L pathway can also limit host cell translation in response to dsRNA stimulation or viral infection. However, while activation of PKR signalling promotes SG assembly, RNase L signalling inhibits assembly of chemically-induced SGs and actively promotes SG disassembly (Burke et al., 2020; Burke et al., 2019). The activation of RNase L results in a translational shut-off that is independent of PKR and phospho-eIF2-alpha and the assembly of small SG-like punctate termed RNase L-bodies (RLB) (Burke et al., 2020). Importantly, while RLBs are sensitive to CHX treatment they do not require G3BP1 for their assembly, and thus are fundamentally distinct from SGs, sharing this property with PGs identified in the current study (Fig. S6). However, these results have been challenged by another study that suggested the specific RNase L ligand 2-5A, and the associated RNase L stimulation, results in the assembly of antiviral stress granules (avSGs) rather than RLBs (Manivannan et al., 2020). In brief, avSGs are different from the RLBs in that they require G3BP1 and activate PKR (Manivannan et al., 2020). These avSGs contains several components of the innate immune pathways including RIG-I, PKR, OAS and RNAse L, and G3BP1 interacts with RIG-I and PKR. Moreover, their assembly is required for IRF3-mediated production and conversely the absence of avSGs sensitized the cells to infection. Therefore, it has proposed that avSGs provide a platform for efficient interaction of RNA ligands and PRR to help mount the antiviral state. In light of these results, we compared our list of PG resident proteins to the previously identified RLB proteome (Burke et al., 2020). This revealed that although PG and RLB share their lack of requirement for G3BP1, they contain different proteins are thus likely to fulfil distinct cellular functions (Fig. S6).

During infection by viruses, several viral products can act as PAMPs recognized by PPRs to amplify the interferon response and create an antiviral state (Kawai and Akira, 2006). Among these, the protein kinase R (PKR) is a major player in antiviral innate immunity. PKR is activated upon binding to double-stranded RNA (dsRNA) such as the genomes of dsRNA viruses, hairpin structures present in ssRNA genomes, or viral replication intermediates (Balachandran et al., 2000). PKR activation triggers the ISR, resulting in global translation inhibition and SG formation. In addition, PKR also acts through innate immune signalling pathways such as nuclear factor-κB (NF-κB), leading to the production of pro-inflammatory cytokines. Therefore, viruses have evolved various strategies to suppress PKR activation and thereby block the ISR and SG formation. These strategies include PKR degradation, PKR inhibition by viral proteins or RNAs, and shielding of dsRNA by viral proteins (Eiermann et al., 2020; Jan et al., 2016). Moreover, many viruses counteract SG formation directly, suggesting that SGs themselves and not only the ISR, are the target. These strategies include encoding proteases that cleave G3BP1, sequestration of SG resident proteins by viral proteins or viral RNAs, or repurposing of stress granules proteins for viral replication (Gaete-Argel et al., 2019; McCormick and Khaperskyy, 2017; Poblete-Duran et al., 2016). A large body of evidence, including functional analysis of virus deletion mutants that abrogate SG formation, or SG composition, suggests that SGs themselves contribute to antiviral response. In particular, the ability of G3BP1 to inhibit enterovirus replication is directly linked to SG formation (Reineke and Lloyd, 2015). Further studies proposed that this is mediated by PKR recruitment to SGs by G3BP1 which potentiates PKR and impair viral replication (Reineke and Lloyd, 2015). Further support to an antiviral role for SGs in amplifying PKR signalling is provided by reduced PKR activation followed pharmacological or genetic disruption of SG formation (Burgess and Mohr, 2018; Manivannan et al., 2020; Ng et al., 2013; Oh et al., 2016; Onomoto et al., 2012; Reineke et al., 2015; Reineke and Lloyd, 2015).

In addition to PKR, several antiviral proteins, including PKR, RIG-I, MDA5, OAS, RNAse L, Trim5, ADAR1, ZAP, cGAS and several RNA helicases can be recruited to SGs during infection, proposing these are antiviral SGs (avSGs) with a role in antiviral signalling (Onomoto et al., 2012; Thulasi Raman et al., 2016; Yoo et al., 2014). These avSGs are essential for IFN signalling in response to several (-) ssRNA viruses as their loss or inhibition resulted in increased replication and impaired antiviral response. The colocalization of viral RNAs and PRRs have suggested that these provide a platform for non-self-sensing and prime interferon production (Ng et al., 2013; Oh et al., 2016; Yoshida et al., 2015). Moreover, the antiviral activity of the zinc-finger antiviral protein (ZAP) correlates with its recruitment to SGs, inhibiting the replication of several viruses through exosome-mediated viral RNA degradation (Law et al., 2019; Leung et al., 2011). Overall, this evidence clearly implicates SGs in antiviral signalling, although specific roles may be restricted to specific cellular contexts or viruses. Our own study implicates the formation of PGs following paracrine signalling by FCV-infected cells in antiviral activity given our observation that PG assembly is associated with impaired viral replication. Our proteomic analysis of PG composition did not detect the accumulation of specific PPR or RLR within PGs. Furthermore, PG-resident proteins clustered into functional categories related to stress like senescence and BRAF signalling, paracrine pathways such as FGFR, DNA damage and rRNA processing. In addition, the analysis of the overall transcriptional reprogramming associated with PG assembly did not highlight typical IFN responses and most differentially expressed genes were functionally clustered in categories related to oxidative phosphorylation, mitochondrial and translation-like ribosome components. Therefore, we speculate that PGs may restrict viral replication through distinct mechanisms than the non-self-sensing platform hypothesis proposed above for SGs and avSGs. Furthermore, it is currently unclear whether infection with other viruses can also induce stimulate the formation of PGs. Interestingly our own studies of SG assembly during infection by the related Murine Norovirus revealed a repurposing of G3BP1 and other SG proteins into viral replication complex rather than PGs assembly (Brocard et al., 2020; Hosmillo et al., 2019).

To date there are only limited examples of paracrine inducers of SG assembly. Angiogenin, an angiogenic factor released by tumour cells which acts as a secreted stress-activated ribonuclease, producing tRNA-derived stress-induced RNAs (tiRNAs) (Emara et al., 2010). These tiRNAs can then be recognized as PKR ligands, promoting translational repression and inducing stress granule assembly (Emara et al., 2010). In addition to this example, prostaglandins are secreted neuroinflammatory molecules, produced after environmental insults to the brain and associated with many neurodegenerative disease (Figueiredo-Pereira et al., 2014). Among these, 15-deoxy-Δ12,14-prostaglandin J2 (15-d-PGJ2) is the most reactive prostaglandin and can covalently modify eIF4A, and cause eIF2α phosphorylation to block translation initiation (Campo et al., 2002; Kim et al., 2007; Marcone and Fitzgerald, 2013). As a result, 15-d-PGJ2 is an endogenously produced trigger of SG assembly and activates the ISR in a cell-nonautonomous manner (Tauber and Parker, 2019). However, preliminary experiments could not demonstrate that VFS stimulation is associated with the production of angiogenin or 15-d-PGJ2 (data not shown). Therefore, we do not rule out several possible origins for the paracrine messenger: nucleic acid (RNA fragment of viral origin, cellular miRNAs), other proteins (cytokines or signalling proteins) or more complexes structures such as exosome particles, which can deliver antiviral sensors to uninfected cells. Assessing these possibilities should be the focus of further studies. Overall, our work expands our understanding of the role of biocondensates in eukaryotes, describing a novel type of cytoplasmic RNP granules formed in response to viral infection and associated with control of viral replication.

## Material and methods

### Cells and viruses

Crandell-Rees feline kidney (CRFK) cells (European Collection of Cell Cultures [ECACC]) were grown in minimum essential medium (MEM, Gibco #31095-029), L-glutamine (Gibco), 1% nonessential amino acids (Gibco) and supplemented with 10% fetal bovine serum (FBS, Gibco), and 1% penicillin-streptomycin in a 5% CO_2_ environment. U2OS cells were grown in DMEM (Dulbecco’s Modified Eagle’s Medium, Gibco #41966), supplemented with 10% (v/v) foetal bovine serum (FBS, Gibco), glutamine and 1% penicillin/streptomycin (Gibco) in a 5% CO_2_ chamber. GFP-G3BP1 expressing and G3BP1 U2OS have been described elsewhere (Burke et al., 2020). Arsenate, puromycin and cycloheximide were all purchased from Sigma. All inhibitors were added to the cell culture media and incubated at 37°C for the indicated times. For viral infections, the FCV strain Urbana was used as previously described (Humoud et al., 2016). The infection titter was determinate by TCID50 is determine which apply serial dilutions to replicate culture of the virus sample. For a multiplicity of infection (MOI) equal to 1 and 0.2 MOI for the virus free supernatant generation. After 1 hour the inoculation was removed, and the cells were incubated with fresh medium for 5 hours post infection (h.p.i.).

### Virus free supernatant preparation

CRFK cells were seeded in 10cm dishes at concentration of 2 × 10^6^ cells and incubated overnight. On the following day, the cells were infected with FCV at 0.2 MOI for 1h and the supernatant was collected 5h.p.i. and then transferred to falcon tube and spun at 4000 rpm at 4°C (Micro 22R, Hettich) for 30 min. After centrifugation, supernatant was filtered through a 0.2 μm Millex filter. The FCV particles were removed via precipitation adding 0.2 M of solid NaCl and 10% of PEG3350 and incubated overnight at 4°C in constant rotation. Then the samples were first centrifuged for 60 minutes at 14000 rpm (Micro 22R, Hettich) at 4°C and then centrifuged at 40000 rpm for 4h30 at 4°C using SW41Ti rotor (Beckman). The supernatant was then UV inactivated at a wavelength of 254 nm for 4 min (3 times) using a cross linker (Stratalinker® UV Crosslinker), filtered through a 0.2 μm Millex filter and stored in -20°C until use.

### Paracrine Granules isolation

For the PGs isolation we follow the protocol previously described in (Brocard et al., 2020), here briefly: G3BP1-GFP-U20S cells were grown O/N on15 cm dishes 80-90% confluent and incubated with 8ml of virus free supernatant for 1h, then washed twice with warm DMEM and scraped out and spun down at 1,500 g 10 min. The cell pellet was either snap freeze or resuspended in 1 ml of lysis buffer (50 mM Tris HCl pH 7.4, 100 mM potassium acetate, 2mM magnesium acetate, 0.5 mM DTT, 50 μg/ml heparin, 0.5% NP40, 1 complete mini EDTA free protease inhibitor tablet/ 50 ml of lysis buffer) and syringe lysis suing 25G 5/8 needle on ice seven time and spun at 300 g, for 5 min at 4°C. The supernatant was transferred and centrifuged again at 18000 g for 20 min at 4C. The supernatant was discarded the pellet was resuspend pellet in 1 ml of SG lysis buffer and spun again at 18000 g for 20 min at 4°C. The pellet was resuspended in 300 μl lysis buffer (granule-enriched fraction). The IP was followed, we performed a pre-clearing with the 60 μl of protein A Dynabeads and the “granule-enriched fraction” for 20-30 min in rotation at 4°C. The supernatant was then incubated 0.5 μg of anti-GFP Ab in rotation at 4°C O/N. Then was added 500 μl of SG lysis buffer to each sample and spun down at 18000g for 20 min at 4°C and the pellet resuspended in 500 μl of SG buffer and 33 μl of Dynabeads and incubated for 3 hrs at 4°C in rotation. The beads were washed once with 1 ml of wash buffer 1 (SG lysis buffer + 2 M Urea) at 4°C for 2 min. Then washed for 5min in 1ml of buffer 2 (SG lysis buffer + 300 mM potassium acetate) at 4°C, 5 min in SG lysis Buffer at 4°C and then seven times with 1ml of TE buffer (10 mM Tris, 1 mM EDTA). The resuspended beads were sent in TE (10 mM Tris HCl and 1 mM EDTA pH 8) buffer for the MS/MS analysis to the Proteomic Facility at university of Bristol.

### Sample preparation for LC-MS/MS analysis

Samples from immunoprecipitation were resuspended in 0.1 M ammonium bicarbonate (ABC), 0.1% sodium deoxycholate, reduced and alkylated using 5 mM TCEP 20 mM chloroacetamide at 70°C for 15 min in darkness. Samples were, then trypsinized using 0.25 µg of sequencing grade modified trypsin (Promega) at 42°C for 4 hours. Sodium deoxylcholate was removed by phase-transfer to ethy lacetate. The resulting tryptic peptides were desalted using in-house StageTips with 3 M Empore SDB-RPS membrane, and dried using vacuum centrifugation. The peptides were reconstituted in 15 µL of Buffer A (0.1% formic acid in water), of which 5 µL was subjected to LC-MS/MS analysis.

### LC-MS/MS analysis

The tryptic peptides were resolved using a Waters nanoACQUITY UPLC system in a single pump trap mode. The peptides were loaded onto a nanoACQUITY 2G-V/MTrap 5µm Symmetry C18 column (180 µm × 20 mm) with 99.5% Buffer A and 0.5% Buffer B (0.1% formic acid in acetonitrile) at 15 µL/min for 3 min. The trapped peptides were eluted and resolved on a BEH C18 column (130 Å, 1.7 µm × 75 µm × 250 mm) using gradients of 3 to 5% B (0-3 min), 8 to 28% B (3-145 min), 28 to 40% B (145-150 min) at 0.3 µL/min. MS/MS was performed on a LTQ Orbitrap Velos mass spectrometer, scanning precursor ions between 400 and 1800 m/z (1 × 106 ions, 60,000 resolution) and selecting the 10 most intense ions for MS/MS with 180 s dynamic exclusion, 10 ppm exclusion width, repeat count = 1, and 30 s repeat duration. Ions with unassigned charge state and MH+1 were excluded from the MS/MS. Maximal ion injection times were 10 ms for FT (one microscan) and 100 ms for LTQ, and the AGC was 1 × 104. The normalized collision energy was 35% with activation Q 0.25 for 10 ms.

### Mass spectrometry data analysis

MaxQuant/Andromeda (version 1.5.2.8) was used to process raw files from LTQ-orbitrap, and search the peak lists against database consist of Uniprot Human proteome (total 71,803 entries, downloaded at 1/12/2018). The search allowed trypsin specificity with maximum two missed-cleavage, and set carbamidomethyl modification on cysteine as a fixed modification and protein N-terminal acetylation and oxidation on methionine as variable modifications. MaxQuant used 4.5 ppm main search tolerance for precursor ions, 0.5 Da MS/MS match tolerance, searching top 8 peaks per 100 Da. False discovery rates for both protein and peptide were 0.01 with minimum seven amino acid peptide length. A label-free quantification was enabled with minimum 2 LFQ ratio counts and a fast LFQ option.

All raw data were analysed with MaxQuant software Two or more unique peptides were used for protein identification and a ratio count of two or more for label free protein quantification in all of the samples. The LFQ intensities were normalised such that at each condition/time point the LFQ intensity values added up to exactly 1,000,000, therefore each protein group value can be regarded as a normalized microshare (performed separately for each sample for all proteins that were present in that sample). The mass spectrometry proteomics data have been deposited to the ProteomeXchange Consortium via the PRIDE [X] partner repository with the dataset identifier PXD021576

### RNA sequencing and data analysis

The granule enriched fraction isolation was isolated as described above, here briefly: G3BP1-GFP-U20S cells were grown O/N on15cm dishes 80-90% confluent and incubated with 8ml of virus free supernatant for 1h, then washed twice with warm DMEM and scraped out and spun down at 1,500 g 10 min. The cell pellet was either snap freeze or resuspended in 1ml of lysis buffer (50 mM Tris HCl pH 7.4, 100 mM potassium acetate, 2 mM magnesium acetate, 0.5 mM DTT, 0.5% NP40, 1 complete mini EDTA free protease inhibitor tablet/ 50 ml of lysis buffer) and syringe lysis using 25G 5/8 needle on ice seven time and spun at 300 g, for 5 min at 4°C. The supernatant was transferred and centrifuged again at18000g for 20min at 4C. The supernatant was discarded the pellet was resuspend pellet in 1 ml of SG lysis buffer and spun again at 18000 g for 2min at 4°C. The pellet was resuspended in 300 μl lysis buffer (granule-enriched fraction). The RNA was extracted with Trizol following the manufactural procedure Invitrogen).

### Library and Sequencing

The precipitated RNA was performed by Novogene to constitute the cDNA library using oligo(dT) beads, then was then fragmented randomly in fragmentation buffer, followed by cDNA synthesis using random hexamers and reverse transcriptase. After first-strand synthesis, a custom second-strand synthesis buffer (Illumina) is added with dNTPs, RNase H and Escherichia coli polymerase I to generate the second strand by nick-translation. The final cDNA library is ready after a round of purification, terminal repair, A-tailing, ligation of sequencing adapters, size selection and PCR enrichment. The library concentration was first quantified using a Qubit 2.0 fluorometer (Life Technologies), and then diluted to 1 ng/μl before checking insert size on an Agilent 2100 and quantifying by quantitative PCR (Q-PCR) (library activity >2 nM). Libraries were fed into Illumina machines according to activity and expected data volume.

### Pre-processing

Quality checks were performed via FastQC (version 0.11.4) (http://www.bioinformatics.babraham.ac.uk/projects/fastqc). Trimmomatic tool (version 0.32) (Bolger et al., 2014) was used for quality trimming and clipping of adapters and repeated sequences. Reads were mapped to the human genome annotation (Gencode Human GRCh38.p12 assembly genome and comprehensive gene annotation, http://www.gencodegenes.org) using the RNA-seq aligner STAR (version 2.5.5b) (Dobin et al., 2013). The function featureCounts from the R package Rsubread (version 1.30.9) (Liao et al., 2014) was used to assign mapped sequencing reads to genomic features. Genomic features were defined by the tool’s in-built NCBI RefSeq annotations for the hg38 genome. R package org.Hs.eg.db (version 3.6.0) [Marc Carlson (2016) org.Mm.eg.db: Genome wide annotation for Mouse. R package version 3.6.0.], Ensembl (accessed March2019 via the R package biomaRt) and GenBank (accessed March 2019 via the R package Annotate) were used to annotate the genomic features.

### Differential expression analyses

Differential expression was performed using the R Bioconductor package EdgeR (version 3.22.5) (Robinson et al., 2010). Filtering of lowly expressed genes was performed, independently for each pairwise comparison, by keeping gens with at least [Equation] counts per million (CPM) in at least 50% of all samples involved in the comparison. EdgeR’s default normalization was applied. CPM values were fitted to a negative binomial generalised log-linear model (GLM) using empirical Bayes tagwise dispersions to estimate the dispersion parameter for each gene. Differential expression was identified using GLM likelihood ratio tests. A paired designed was used when comparing SGs vs Total in treatment and when comparing SGs vs Total in Control, and a two groups design was used when comparing SGs Treatment vs Control and when comparing Total Treatment vs Control.

### Immunofluorescence (IF) microscopy

Cells in 24-well plate were washed with pre-warmed DPBS and immediately incubated in 500 µl of fixing solution (4% PFA in PBS) for 15 min at room temperature (RT), and further permeabilised with 500 µl of 0.1% Triton X-100 in PBS for 10 min at RT. For the N6A Methyladenosine hybridisation only, the fixation was performed adding 200 µl per well of ice-cold methanol, the plate was then incubated at 20°C for 10 minutes, and then washes with PBS. Blocking was carried out with 1 ml of blocking solution (1% BSA in PBS) for 1 h at RT. Fixed cells were then incubated with primary antibody in 0.5% BSA PBS for 2 h at room temperature (RT), and washed three times with PBS prior to the addition of a secondary antibodies for 1 h at RT in the dark. Finally, cells were washed three times with PBS and mounted on the slide with ProLong Diamond antifade with 4′,6-diamidino-2-phenylindole (DAPI; Life Technologies, #P36966). Confocal images were acquired on a Nikon Ti-Eclipse A1M microscope fitted with a 60× oil immersion objective using 488 nm, 561 nm and 405 nm laser excitation lines.

### Preparation of RNA samples and RT-qPCR

Total RNA was extracted by Trizol (Invitrogen) following the manufacturing procedures and then subjected to Reverse Transcription (primer design) with oligo(dT)15 and random primers following the manufacturer’s protocol. Subsequently, the real time PCR was performed using specific primers: The samples were analysed in triplicate with SYBR GREEN dye (Primer Design Precision Master mix) on an ABI StepOnePlus quantitative PCR instrument (Applied Biosystems). The comparative Ct method was employed to measure amplification of specific mRNAs vs. the total level of β-actin or to tubulin where indicated.

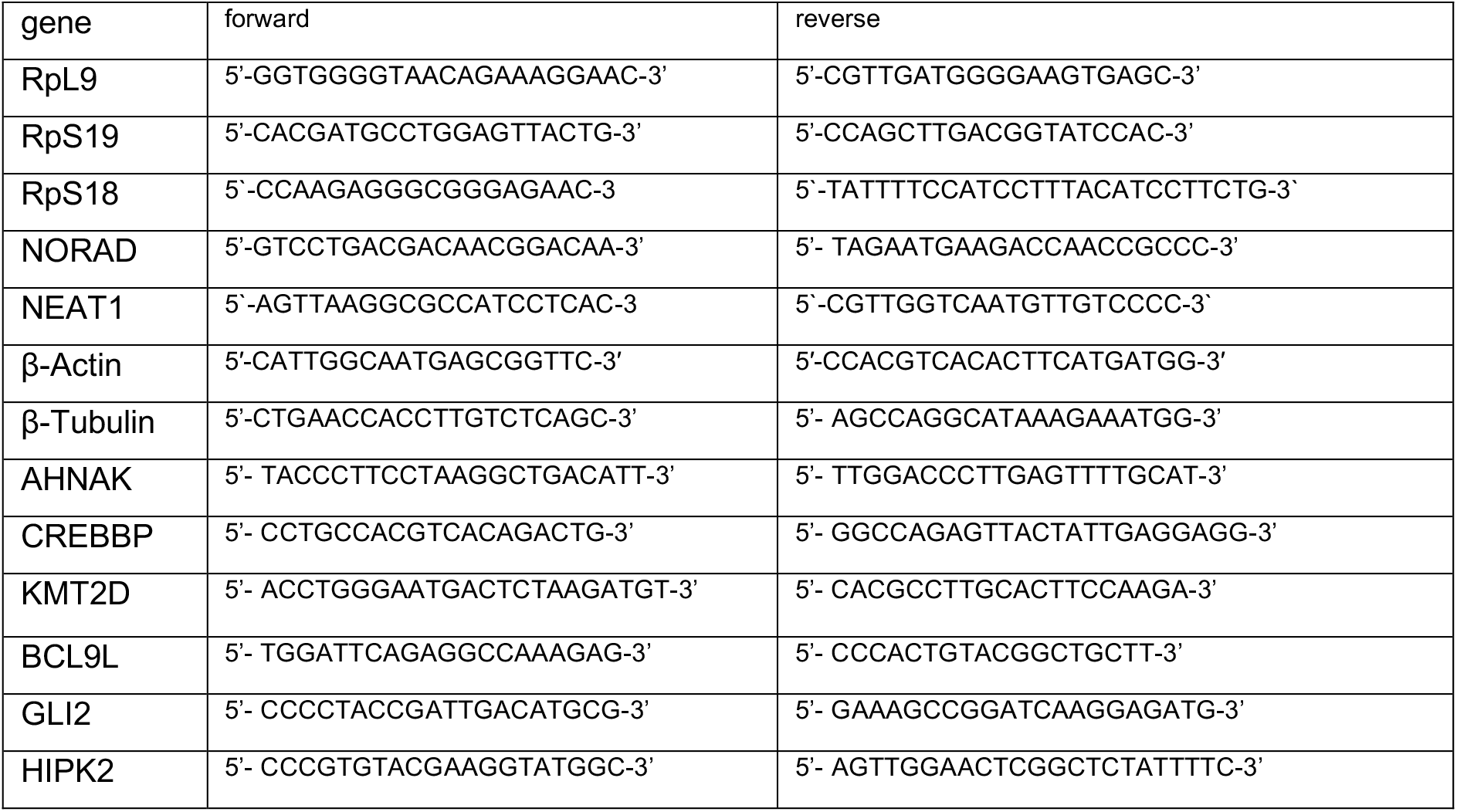

### Western Blotting

To prepare total cell extracts, GFP U2OS cells were washed twice with cold phosphate-buffered saline (PBS) and then lysed in buffer High salt buffer (50 mM Tris HCl pH 7.5, 350 mM NaCl, 1 mM MgCl2, 0.5 mM EDTA, 0.1 mM EGTA and 1% (v/v) Triton X-100), protease inhibitor cocktail was added to the lysis buffer just before use. Lysates were incubated on ice for 30 minutes, then centrifuged at 4°C, at maximum speed for 10 min and then the supernatant was collected. Protein concentrations were determined by Bradford reagent (Bio-Rad). The lysed was heated at 95°C for 5 min in sample buffer (62.5 mM Tris-HCl, 7% (w/v) SDS, 20% (w/v) sucrose and 0.01% (w/v) bromophenol blue) and subjected to polyacrylamide gel electrophoresis and electrophoretic transfer to nitrocellulose (for eIF2alpha blot) or PVDF membranes. Membranes were then blocked in tris-buffered saline (TBS) Tween 20 containing 5% (w/v) skimmed milk powder for 30 min at room temperature. The membranes were probed with the primary antibody indicated below in 3% BSA overnight at 4°C, followed by incubation with the appropriate peroxidase labelled secondary antibodies (Dako) and chemiluminescence development using the Clarity Western ECL Substrate (Bio-Rad). The results were acquired using the VILBER imaging system.

### Antibodies

| Antibody | Company | Catalogue number | Dilution | WB/IF |
| --- | --- | --- | --- | --- |
| HRP-coupled goat anti-mouse | Dako | P0447 | 1:1000 to 1:5000 | WB |
| HRP-coupled goat anti-rabbit | Dako | P0448 | 1:1000 to 1:5000 | WB |
| eIF2α | Cell Signaling Technology | #9722 | 1:1000 | WB |
| eIF2alpha Ser51 | Cell Signaling Technology | #9721 | 1:1000 | WB |
| eIF3b | Santa Cruz | Sc-16377 | 1:600 | IF |
| eIF4E | Cell signaling | #9742 | 1:1000 | WB |
| p38 | Cell Signaling Technology | #9212 | 1:1000 | WB |
| p38 T180/Y182 | Cell Signaling Technology | #9211 | 1:1000 | WB |
| mTOR | Cell Signaling Technology | #9283 | 1:1000 | WB |
| mTOR Ser2448 | Cell Signaling Technology | #5536 | 1:1000 | WB |
| ERK1/2 | Cell Signaling Technology | #9102 | 1:1000 | WB |
| ERK1/2 Thr202/Tyr204 | Cell Signaling Technology | #9101 | 1:1000 | WB |
| AKT | Cell Signaling Technology | #9272 | 1:1000 | WB |
| AKT S473 | Cell Signaling Technology | #4060 | 1:1000 | WB |
| ELAVL1 | Santa Cruz | Sc-5261 | 1:2000<br>1:300 | WB<br>IF |
| eIF3b | Santa Cruz | Sc-16377 | 1:600 | IF |
| G3BP1 | BD bioscience | #611127 | 1:1000<br>1:600 | WB<br>IF |
| HNRNPK | Protein tech | #11426-1-AP | 1:600 | IF |
| DSG1 | Protein tech | #24587-1-AP | 1:600 | IF |
| ERH | Protein tech | #15974-1-AP | 1:600 | IF |
| FUS | Protein tech | #11570-1-AP | 1:600 | IF |
| SFRS7 | Novus bio | NBP1-92382 | 1:300 | IF |
| FXR1 | Merk Millipore | #05-1529 | 1:600 | IF |
| FXR1 | Novus Bio | NBP1-89546 | 1:600 | IF |
| RBMX | Sigma | HPA057707 | 1:600 | IF |
| THRAP3 | Protein tech | #19744-1-AP | 1:600 | IF |
| RNPS1 | Protein tech | #10555-1-AP | 1:600 | IF |
| NOLC1 | Protein tech | #11815-1-AP | 1:600<br>1:1000 | IF<br>WB |
| YTHDF3 | Protein tech | #25537-1-AP | 1:600<br>1:1000 | IF<br>WB |
| UBAP2L | Abcam | AB138309 | 1:300 | IF |
| B23 | SIGMA | B0556 | 1:600 | IF |
| N6A Methyladenosine | Merk | ABE572 | 1:300 | IF |
| Lamin B | Protein tech | #12987-1-AP | 1:1000 | WB |
| β-actin | Protein tech | #66009-1-AP | 1:5000 | WB |
| GAPDH | Santa Cruz | Sc-32233 | 1:20000 | WB |
| Tubulin | Santa Cruz | Sc-5546 | 1:2000 | WB |
| NS7 | Described in (Humoud et al., 2016) |  | 1:2000 | WB |
| Alexa-Fluor 555 anti-rabbit | Invitrogen | A21428 | 1:600 | IF |
| Alexa-Fluor 555 anti-mouse | Invitrogen | A21424 | 1:600 | IF |
| Alexa-Fluor 555 anti-goat | Invitrogen | A21432 | 1:600 | IF |
| Alexa-Fluor 647 anti-mouse | Invitrogen | A21463 | 1:600 | IF |

### FRAP and time-lapse

GFP-G3BP1 U2OS cells were cultured on glass bottom on 24-well plate the day before. The following day the cells were treated either with the virus-free supernatant or with arsenite for 1h at 37°C before to be subject to FRAP. The cells were imaged using Nikon AR1 confocal microscope and maintained at 37°C, 5% CO2 during imaging acquisition. FRAP experiments were performed as follows: the region of interest (ROI) was defined about the size of the granule (∼2 μm) for bleaching and acquisition area (BL), then we determined a similar ROI size outside of the cells, considered as a region of noise (BG) and finally the last ROI was defined around a granule used as a reference for the intensity (REF) to correct the beaching during the imaging acquisition. The bleaching was carried out with full laser power for 30 sec and the acquisition of each ROIs was set every 3 seconds. The background noise was subtracted from each given time point (**BL_corr1(t)** = BL(t) – BG(t) and from the **REF_corr1(t)** = REF(t) – BG(t)), then in order to calculate the normalize corrected bleach value the following formula has applied: **BL_corr2(t)** = BL_corr1(t) / REF_corr1(t) = [BL(t) – BG(t)] / [REF(t)-BG(t)]. Finally normalise to the mean of the pre-bleach intensity **BL_corr3(t)** = BL_corr2(t) / BL_corr2(pre-bleach). The curve for the exponential GFP recovery was determined using GraphPad Prims8.

### Ribopuromycylation assay

Quantification of de novo protein synthesis was performed as described in (Brocard et al., 2020). U2OS cells were treated with 10 μg/ml of puromycin (Sigma), for 5 minutes at 37°C, and then with 180 μM of emetin (Sigma) for 2 minutes at 37°C, before fixation were washed three times with pre-warmed DMEM and fixed with 4% PFA in PBS for 10 minutes at RT. Fluorescence intensities were quantified by using Image J software package Fiji.s

## Acknowledgements

The authors thank Alessia Ruggieri (University of Heidelberg) for providing critical comments on the study and manuscript.

## Data availability

The raw RNA-seq data and mapping results can be found as part of GEO repository database, submission GSE171655. The mass spectrometry proteomics data have been deposited to the ProteomeXchange Consortium via the PRIDE partner repository with the dataset identifier PXD021881.

## References

An, H., J.T. Tan, and T.A. Shelkovnikova. 2019. Stress granules regulate stress-induced paraspeckle assembly. J Cell Biol. 218:4127–4140.

Anders, M., I. Chelysheva, I. Goebel, T. Trenkner, J. Zhou, Y. Mao, S. Verzini, S.B. Qian, and Z. Ignatova. 2018. Dynamic m(6)A methylation facilitates mRNA triaging to stress granules. Life Sci Alliance. 1:e201800113.

Anderson, P., N. Kedersha, and P. Ivanov. 2015. Stress granules, P-bodies and cancer. Biochim Biophys Acta. 1849:861–870.

Aulas, A., M.M. Fay, S.M. Lyons, C.A. Achorn, N. Kedersha, P. Anderson, and P. Ivanov. 2017. Stress-specific differences in assembly and composition of stress granules and related foci. J Cell Sci. 130:927–937.

Bailey, T.L., M. Boden, F.A. Buske, M. Frith, C.E. Grant, L. Clementi, J. Ren, W.W. Li, and W.S. Noble. 2009. MEME SUITE: tools for motif discovery and searching. Nucleic Acids Res. 37:W202–208.

Balachandran, S., P.C. Roberts, L.E. Brown, H. Truong, A.K. Pattnaik, D.R. Archer, and G.N. Barber. 2000. Essential role for the dsRNA-dependent protein kinase PKR in innate immunity to viral infection. Immunity. 13:129–141.

Bolger, A.M., M. Lohse, and B. Usadel. 2014. Trimmomatic: a flexible trimmer for Illumina sequence data. Bioinformatics. 30:2114–2120.

Brocard, M., V. Iadevaia, P. Klein, B. Hall, G. Lewis, J. Lu, J. Burke, M.M. Willcocks, R. Parker, I.G. Goodfellow, A. Ruggieri, and N. Locker. 2020. Norovirus infection results in eIF2alpha independent host translation shut-off and remodels the G3BP1 interactome evading stress granule formation. PLoS Pathog. 16:e1008250.

Burgess, H.M., and I. Mohr. 2018. Defining the Role of Stress Granules in Innate Immune Suppression by the Herpes Simplex Virus 1 Endoribonuclease VHS. J Virol. 92.

Burke, J.M., E.T. Lester, D. Tauber, and R. Parker. 2020. RNase L promotes the formation of unique ribonucleoprotein granules distinct from stress granules. J Biol Chem. 295:1426–1438.

Burke, J.M., S.L. Moon, T. Matheny, and R. Parker. 2019. RNase L Reprograms Translation by Widespread mRNA Turnover Escaped by Antiviral mRNAs. Mol Cell. 75:1203–1217 e1205.

Campo, P.A., S. Das, C.H. Hsiang, T. Bui, C.E. Samuel, and D.S. Straus. 2002. Translational regulation of cyclin D1 by 15-deoxy-delta(12,14)-prostaglandin J(2). Cell Growth Differ. 13:409–420.

Cirillo, L., A. Cieren, S. Barbieri, A. Khong, F. Schwager, R. Parker, and M. Gotta. 2020. UBAP2L Forms Distinct Cores that Act in Nucleating Stress Granules Upstream of G3BP1. Curr Biol. 30:698–707 e696.

Corbet, G.A., and R. Parker. 2019. RNP Granule Formation: Lessons from P-Bodies and Stress Granules. Cold Spring Harb Symp Quant Biol. 84:203–215.

Dobin, A., C.A. Davis, F. Schlesinger, J. Drenkow, C. Zaleski, S. Jha, P. Batut, M. Chaisson, and T.R. Gingeras. 2013. STAR: ultrafast universal RNA-seq aligner. Bioinformatics. 29:15–21.

Edupuganti, R.R., S. Geiger, R.G.H. Lindeboom, H. Shi, P.J. Hsu, Z. Lu, S.Y. Wang, M.P.A. Baltissen, P. Jansen, M. Rossa, M. Muller, H.G. Stunnenberg, C. He, T. Carell, and M. Vermeulen. 2017. N(6)-methyladenosine (m(6)A) recruits and repels proteins to regulate mRNA homeostasis. Nat Struct Mol Biol. 24:870–878.

Eiermann, N., K. Haneke, Z. Sun, G. Stoecklin, and A. Ruggieri. 2020. Dance with the Devil: Stress Granules and Signaling in Antiviral Responses. Viruses. 12.

Elbaum-Garfinkle, S. 2019. Matter over mind: Liquid phase separation and neurodegeneration. J Biol Chem. 294:7160–7168.

Emara, M.M., P. Ivanov, T. Hickman, N. Dawra, S. Tisdale, N. Kedersha, G.F. Hu, and P. Anderson. 2010. Angiogenin-induced tRNA-derived stress-induced RNAs promote stress-induced stress granule assembly. J Biol Chem. 285:10959–10968.

Figueiredo-Pereira, M.E., P. Rockwell, T. Schmidt-Glenewinkel, and P. Serrano. 2014. Neuroinflammation and J2 prostaglandins: linking impairment of the ubiquitin-proteasome pathway and mitochondria to neurodegeneration. Front Mol Neurosci. 7:104.

Gaete-Argel, A., C.L. Marquez, G.P. Barriga, R. Soto-Rifo, and F. Valiente-Echeverria. 2019. Strategies for Success. Viral Infections and Membraneless Organelles. Front Cell Infect Microbiol. 9:336.

Guillen-Boixet, J., A. Kopach, A.S. Holehouse, S. Wittmann, M. Jahnel, R. Schlussler, K. Kim, I. Trussina, J. Wang, D. Mateju, I. Poser, S. Maharana, M. Ruer-Gruss, D. Richter, X. Zhang, Y.T. Chang, J. Guck, A. Honigmann, J. Mahamid, A.A. Hyman, R.V. Pappu, S. Alberti, and T.M. Franzmann. 2020. RNA-Induced Conformational Switching and Clustering of G3BP Drive Stress Granule Assembly by Condensation. Cell. 181:346–361 e317.

Hofmann, S., V. Cherkasova, P. Bankhead, B. Bukau, and G. Stoecklin. 2012. Translation suppression promotes stress granule formation and cell survival in response to cold shock. Mol Biol Cell. 23:3786–3800.

Hofmann, S., N. Kedersha, P. Anderson, and P. Ivanov. 2020. Molecular mechanisms of stress granule assembly and disassembly. Biochim Biophys Acta Mol Cell Res:118876.

Hosmillo, M., J. Lu, M.R. McAllaster, J.B. Eaglesham, X. Wang, E. Emmott, P. Domingues, Y. Chaudhry, T.J. Fitzmaurice, M.K. Tung, M.D. Panas, G. McInerney, N. Locker, C.B. Wilen, and I.G. Goodfellow. 2019. Noroviruses subvert the core stress granule component G3BP1 to promote viral VPg-dependent translation. Elife. 8.

Hubstenberger, A., M. Courel, M. Benard, S. Souquere, M. Ernoult-Lange, R. Chouaib, Z. Yi, J.B. Morlot, A. Munier, M. Fradet, M. Daunesse, E. Bertrand, G. Pierron, J. Mozziconacci, M. Kress, and D. Weil. 2017. P-Body Purification Reveals the Condensation of Repressed mRNA Regulons. Mol Cell. 68:144–157 e145.

Humoud, M.N., N. Doyle, E. Royall, M.M. Willcocks, F. Sorgeloos, F. van Kuppeveld, L.O. Roberts, I.G. Goodfellow, M.A. Langereis, and N. Locker. 2016. Feline Calicivirus Infection Disrupts Assembly of Cytoplasmic Stress Granules and Induces G3BP1 Cleavage. J Virol. 90:6489–6501.

Jain, S., J.R. Wheeler, R.W. Walters, A. Agrawal, A. Barsic, and R. Parker. 2016. ATPase-Modulated Stress Granules Contain a Diverse Proteome and Substructure. Cell. 164:487–498.

Jan, E., I. Mohr, and D. Walsh. 2016. A Cap-to-Tail Guide to mRNA Translation Strategies in Virus-Infected Cells. Annu Rev Virol. 3:283–307.

Kawai, T., and S. Akira. 2006. Innate immune recognition of viral infection. Nat Immunol. 7:131–137.

Kedersha, N., M.R. Cho, W. Li, P.W. Yacono, S. Chen, N. Gilks, D.E. Golan, and P. Anderson. 2000. Dynamic shuttling of TIA-1 accompanies the recruitment of mRNA to mammalian stress granules. J Cell Biol. 151:1257–1268.

Khong, A., S. Jain, T. Matheny, J.R. Wheeler, and R. Parker. 2018. Isolation of mammalian stress granule cores for RNA-Seq analysis. Methods. 137:49–54.

Khong, A., T. Matheny, S. Jain, S.F. Mitchell, J.R. Wheeler, and R. Parker. 2017. The Stress Granule Transcriptome Reveals Principles of mRNA Accumulation in Stress Granules. Mol Cell. 68:808–820 e805.

Kim, W.J., J.H. Kim, and S.K. Jang. 2007. Anti-inflammatory lipid mediator 15d-PGJ2 inhibits translation through inactivation of eIF4A. EMBO J. 26:5020–5032.

Law, L.M.J., B.S. Razooky, M.M.H. Li, S. You, A. Jurado, C.M. Rice, and M.R. MacDonald. 2019. ZAP’s stress granule localization is correlated with its antiviral activity and induced by virus replication. PLoS Pathog. 15:e1007798.

Leung, A.K., S. Vyas, J.E. Rood, A. Bhutkar, P.A. Sharp, and P. Chang. 2011. Poly(ADP-ribose) regulates stress responses and microRNA activity in the cytoplasm. Mol Cell. 42:489–499.

Liao, Y., G.K. Smyth, and W. Shi. 2014. featureCounts: an efficient general purpose program for assigning sequence reads to genomic features. Bioinformatics. 30:923–930.

Manivannan, P., M.A. Siddiqui, and K. Malathi. 2020. RNase L Amplifies Interferon Signaling by Inducing Protein Kinase R-Mediated Antiviral Stress Granules. J Virol. 94.

Marcone, S., and D.J. Fitzgerald. 2013. Proteomic identification of the candidate target proteins of 15-deoxy-delta12,14-prostaglandin J2. Proteomics. 13:2135–2139.

Markmiller, S., S. Soltanieh, K.L. Server, R. Mak, W. Jin, M.Y. Fang, E.C. Luo, F. Krach, D. Yang, A. Sen, A. Fulzele, J.M. Wozniak, D.J. Gonzalez, M.W. Kankel, F.B. Gao, E.J. Bennett, E. Lecuyer, and G.W. Yeo. 2018. Context-Dependent and Disease-Specific Diversity in Protein Interactions within Stress Granules. Cell. 172:590–604 e513.

Mateju, D., and J.A. Chao. 2021. Stress granules: regulators or by-products? FEBS J.

Matheny, T., B.S. Rao, and R. Parker. 2019. Transcriptome-Wide Comparison of Stress Granules and P-Bodies Reveals that Translation Plays a Major Role in RNA Partitioning. Mol Cell Biol. 39.

McCormick, C., and D.A. Khaperskyy. 2017. Translation inhibition and stress granules in the antiviral immune response. Nat Rev Immunol. 17:647–660.

Namkoong, S., A. Ho, Y.M. Woo, H. Kwak, and J.H. Lee. 2018. Systematic Characterization of Stress-Induced RNA Granulation. Mol Cell. 70:175–187 e178.

Ng, C.S., M. Jogi, J.S. Yoo, K. Onomoto, S. Koike, T. Iwasaki, M. Yoneyama, H. Kato, and T. Fujita. 2013. Encephalomyocarditis virus disrupts stress granules, the critical platform for triggering antiviral innate immune responses. J Virol. 87:9511–9522.

Oh, S.W., K. Onomoto, M. Wakimoto, K. Onoguchi, F. Ishidate, T. Fujiwara, M. Yoneyama, H. Kato, and T. Fujita. 2016. Leader-Containing Uncapped Viral Transcript Activates RIG-I in Antiviral Stress Granules. PLoS Pathog. 12:e1005444.

Onomoto, K., M. Jogi, J.S. Yoo, R. Narita, S. Morimoto, A. Takemura, S. Sambhara, A. Kawaguchi, S. Osari, K. Nagata, T. Matsumiya, H. Namiki, M. Yoneyama, and T. Fujita. 2012. Critical role of an antiviral stress granule containing RIG-I and PKR in viral detection and innate immunity. PLoS One. 7:e43031.

Peng, Z., M.J. Mizianty, and L. Kurgan. 2014. Genome-scale prediction of proteins with long intrinsically disordered regions. Proteins. 82:145–158.

Poblete-Duran, N., Y. Prades-Perez, J. Vera-Otarola, R. Soto-Rifo, and F. Valiente-Echeverria. 2016. Who Regulates Whom? An Overview of RNA Granules and Viral Infections. Viruses. 8.

Reineke, L.C., N. Kedersha, M.A. Langereis, F.J. van Kuppeveld, and R.E. Lloyd. 2015. Stress granules regulate double-stranded RNA-dependent protein kinase activation through a complex containing G3BP1 and Caprin1. mBio. 6:e02486.

Reineke, L.C., and R.E. Lloyd. 2015. The stress granule protein G3BP1 recruits protein kinase R to promote multiple innate immune antiviral responses. J Virol. 89:2575–2589.

Reineke, L.C., and J.R. Neilson. 2019. Differences between acute and chronic stress granules, and how these differences may impact function in human disease. Biochem Pharmacol. 162:123–131.

Riggs, C.L., N. Kedersha, P. Ivanov, and P. Anderson. 2020. Mammalian stress granules and P bodies at a glance. J Cell Sci. 133.

Robinson, M.D., D.J. McCarthy, and G.K. Smyth. 2010. edgeR: a Bioconductor package for differential expression analysis of digital gene expression data. Bioinformatics. 26:139–140.

Roobol, A., M.J. Carden, R.J. Newsam, and C.M. Smales. 2009. Biochemical insights into the mechanisms central to the response of mammalian cells to cold stress and subsequent rewarming. FEBS J. 276:286–302.

Sanders, D.W., N. Kedersha, D.S.W. Lee, A.R. Strom, V. Drake, J.A. Riback, D. Bracha, J.M. Eeftens, A. Iwanicki, A. Wang, M.T. Wei, G. Whitney, S.M. Lyons, P. Anderson, W.M. Jacobs, P. Ivanov, and C.P. Brangwynne. 2020. Competing Protein-RNA Interaction Networks Control Multiphase Intracellular Organization. Cell. 181:306–324 e328.

Szklarczyk, D., A.L. Gable, D. Lyon, A. Junge, S. Wyder, J. Huerta-Cepas, M. Simonovic, N.T. Doncheva, J.H. Morris, P. Bork, L.J. Jensen, and C.V. Mering. 2019. STRING v11: protein-protein association networks with increased coverage, supporting functional discovery in genome-wide experimental datasets. Nucleic Acids Res. 47:D607–D613.

Taniuchi, S., M. Miyake, K. Tsugawa, M. Oyadomari, and S. Oyadomari. 2016. Integrated stress response of vertebrates is regulated by four eIF2alpha kinases. Sci Rep. 6:32886.

Tauber, D., and R. Parker. 2019. 15-Deoxy-Delta(12,14)-prostaglandin J2 promotes phosphorylation of eukaryotic initiation factor 2alpha and activates the integrated stress response. J Biol Chem. 294:6344–6352.

Tauber, D., G. Tauber, A. Khong, B. Van Treeck, J. Pelletier, and R. Parker. 2020. Modulation of RNA Condensation by the DEAD-Box Protein eIF4A. Cell. 180:411–426 e416.

Thulasi Raman, S.N., G. Liu, H.M. Pyo, Y.C. Cui, F. Xu, L.E. Ayalew, S.K. Tikoo, and Y. Zhou. 2016. DDX3 Interacts with Influenza A Virus NS1 and NP Proteins and Exerts Antiviral Function through Regulation of Stress Granule Formation. J Virol. 90:3661–3675.

Van Treeck, B., D.S.W. Protter, T. Matheny, A. Khong, C.D. Link, and R. Parker. 2018. RNA self-assembly contributes to stress granule formation and defining the stress granule transcriptome. Proc Natl Acad Sci U S A. 115:2734–2739.

Wang, J., J.M. Choi, A.S. Holehouse, H.O. Lee, X. Zhang, M. Jahnel, S. Maharana, R. Lemaitre, A. Pozniakovsky, D. Drechsel, I. Poser, R.V. Pappu, S. Alberti, and A.A. Hyman. 2018. A Molecular Grammar Governing the Driving Forces for Phase Separation of Prion-like RNA Binding Proteins. Cell. 174:688–699 e616.

Wolozin, B., and P. Ivanov. 2019. Stress granules and neurodegeneration. Nat Rev Neurosci. 20:649–666.

Yang, P., C. Mathieu, R.M. Kolaitis, P. Zhang, J. Messing, U. Yurtsever, Z. Yang, J. Wu, Y. Li, Q. Pan, J. Yu, E.W. Martin, T. Mittag, H.J. Kim, and J.P. Taylor. 2020. G3BP1 Is a Tunable Switch that Triggers Phase Separation to Assemble Stress Granules. Cell. 181:325–345 e328.

Yoo, H., C. Triandafillou, and D.A. Drummond. 2019. Cellular sensing by phase separation: Using the process, not just the products. J Biol Chem. 294:7151–7159.

Yoo, J.S., K. Takahasi, C.S. Ng, R. Ouda, K. Onomoto, M. Yoneyama, J.C. Lai, S. Lattmann, Y. Nagamine, T. Matsui, K. Iwabuchi, H. Kato, and T. Fujita. 2014. DHX36 enhances RIG-I signaling by facilitating PKR-mediated antiviral stress granule formation. PLoS Pathog. 10:e1004012.

Yoshida, A., R. Kawabata, T. Honda, K. Tomonaga, T. Sakaguchi, and T. Irie. 2015. IFN-beta-inducing, unusual viral RNA species produced by paramyxovirus infection accumulated into distinct cytoplasmic structures in an RNA-type-dependent manner. Front Microbiol. 6:804.

Youn, J.Y., B.J.A. Dyakov, J. Zhang, J.D.R. Knight, R.M. Vernon, J.D. Forman-Kay, and A.C. Gingras. 2019. Properties of Stress Granule and P-Body Proteomes. Mol Cell. 76:286–294.

Zhou, J., J. Wan, X. Gao, X. Zhang, S.R. Jaffrey, and S.B. Qian. 2015. Dynamic m(6)A mRNA methylation directs translational control of heat shock response. Nature. 526:591–594.

Zhou, Y., B. Zhou, L. Pache, M. Chang, A.H. Khodabakhshi, O. Tanaseichuk, C. Benner, and S.K. Chanda. 2019. Metascape provides a biologist-oriented resource for the analysis of systems-level datasets. Nat Commun. 10:1523.

